# Cell-type-specific transposable element demethylation and TAD remodeling in the aging mouse brain

**DOI:** 10.1101/2025.04.21.648266

**Authors:** Qiurui Zeng, Wei Tian, Amit Klein, Anna Bartlett, Hanqing Liu, Joseph R. Nery, Rosa G. Castanon, Julia Osteen, Nicholas D. Johnson, Wenliang Wang, Wubin Ding, Huaming Chen, Jordan Altshul, Mia Kenworthy, Cynthia Valadon, William Owens, Zhanghao Wu, Maria Luisa Amaral, Yuru Song, Cindy Tatiana Báez-Becerra, Silvia Cho, Chumo Chen, Jackson Willier, Stella Cao, Jonathan Rink, Jasper Lee, Ariana Barcoma, Jessica Arzavala, Nora Emerson, Yuancheng Ryan Lu, Bing Ren, M. Margarita Behrens, Joseph R. Ecker

**Author notes:** Correspondence (M.M.B), (J.R.E.).

## Abstract

Aging is a major risk factor for neurodegenerative diseases, yet underlying epigenetic mechanisms remain unclear. Here, we generated a comprehensive single-nucleus cell atlas of brain aging across multiple brain regions, comprising 132,551 single-cell methylomes and 72,666 joint chromatin conformation-methylome nuclei. Integration with companion transcriptomic and chromatin accessibility data yielded a cross-modality taxonomy of 36 major cell types. We observed that age-related methylation changes were more pronounced in non-neuronal cells. Transposable element methylation alone distinguished age groups, showing cell-type-specific genome-wide demethylation. Chromatin conformation analysis demonstrated age-related increases in TAD boundary strength with enhanced accessibility at CTCF binding sites. Spatial transcriptomics across 895,296 cells revealed regional heterogeneity during aging within identical cell types. Finally, we developed novel deep-learning models that accurately predict age-related gene expression changes using multi-modal epigenetic features, providing mechanistic insights into gene regulation. This dataset advances our understanding of brain aging and offers potential translational applications.

**Highlights:**

- Single-cell multi-omic profiling maps the epigenetic and spatial transcriptomic landscape of brain aging across multiple regions.
- Cell-type-specific genome-wide demethylation of retrotransposable elements correlates with increased chromatin accessibility and expression.
- Elevated TAD boundary strength emerges as a unique marker of brain aging associated with CTCF gaining accessibility.
- A novel deep-learning model reveals the significance of epigenetic features on age-related transcriptomic changes across genes.

## INTRODUCTION

Aging is the primary risk factor for neurodegenerative diseases such as Alzheimer’s, Parkinson’s, Huntington’s, and ALS^1^. Between 2015 and 2050, the proportion of the world’s population over 60 years will nearly double from 12% to 22%^2^. Age-related brain changes, particularly in regions critical for attention, memory, emotion, and motor functions, severely impact life quality^1^. Understanding how aging contributes to the progression of these diseases is essential in implementing preventive strategies and developing targeted therapies.

At the molecular level, hallmarks of aging include chronic inflammation, mitochondrial dysfunction, genome instability, and epigenetic changes^3^. Epigenetic changes have recently been proposed to play a causal and upstream role in physiological aging in mammals^4,5^, and partial epigenetic reprogramming emerges as a promising way to reset these epigenetic changes and achieve functional rejuvenation and lifespan extension^6–8^. Among the epigenetic modifications, cytosine DNA methylation (5mC), a covalent genome modification in post-mitotic cells throughout their lifespan, is associated with neuronal function, behavior, and various diseases. It has been shown that the methylation level of several hundred CpG sites can serve as a reliable estimator of chronological age across mammalian species, commonly referred to as DNAm clock^9,10^. However, it remains challenging to determine whether these age-associated methylation changes reflect biological processes or merely stochastic variations^11^ that increase with age, underscoring the value of generating comprehensive whole-genome methylation data. In addition to 5mC that occurs at CpG sites (mCG) in mammalian genomes, non-CpG cytosine methylation (mCH, where H can be A, C, or T) is also prevalent in neurons^12,13^. Together, CpG and CpH methylation modulate transcription factor (TF) binding and gene transcription through dynamic occurrence at regulatory elements and gene bodies. The local depletion of DNA methylation is a reliable marker of active regulatory elements such as enhancers.

Beyond identifying putative regulatory elements, DNA methylation also serves as a critical epigenetic mechanism for genome stability maintenance, with a key function in silencing transposable elements (TEs). During aging, loss of methylation can derepress TEs such as Long Interspersed Element (LINE) retrotransposons and accelerate aging^14,15^. The derepression of TEs will trigger DNA damage, inflammatory responses, and altered gene expression patterns, ultimately contributing to cellular dysfunction and age-related decline^16–18^.

Besides DNA methylation, cis-regulatory elements in complex mammalian genomes can operate over long distances to regulate target genes. Disruption of genome stability and 3D genome organization by DNA double-strand breaks in neurons are pathological steps in the progression of neurodegeneration during Alzheimer’s disease ^19^. During both human and mouse aging, cerebellar granule neurons establish ultra-long-range intrachromosomal contacts that are rarely seen in young, healthy cells^20^. Global reorganization of the nuclear landscape is observed in senescent cells, with a loss of local interaction in senescence-associated heterochromatin foci formation^21^. Understanding the relationships between the physical contact frequency of enhancers and promoters and their impact on genes’ epigenetics and transcriptomic status is crucial for decoding the molecular alterations during brain aging.

Recent studies have profiled brain-wide cell-type-specific transcriptome changes during aging^22^. Additionally, some studies have examined specific epigenetic changes during normal aging in targeted regions or cell types^20,23–25^. However, there has not been comprehensive research investigating multi-modality epigenetic changes during aging at a multi-brain region level. To fully understand the regulatory mechanisms underlying age-related gene expression changes and the differing vulnerabilities of brain regions during aging, it is essential to capture data from multiple brain regions and employ various epigenomic modalities. In this study, we have developed the most comprehensive single-cell multi-omic brain aging dataset to date. Our dataset spans methylation, chromatin conformation, and spatial transcriptomics, complemented by our companion study (Amaral et al.) that provides chromatin accessibility and transcriptome profiles across eight brain regions vulnerable in neurodegenerative diseases. With this dataset, we present a detailed analysis of age-related epigenetic and transcriptional changes in diverse cell types throughout the brain.

Through analysis with our single-cell whole-genome methylation data, we reveal that non-neuronal cells exhibit more age-related methylation changes than neurons. Our analysis revealed significant activator protein 1 (AP-1) family (for example, Fos, Jun) motif enrichment in neuronal hypomethylated regions, while Sox family transcription factors were enriched in glial hypermethylated regions. Remarkably, we also found that the methylation patterns in transposon regions can not only distinguish cell types but also effectively segregate cells into different age groups. At the cell-type resolution, we observed locus-specific demethylation in TE regions, which correlates with increased chromatin accessibility and RNA expression, highlighting cell types that are particularly affected by TE activation. Insights from our single-cell chromatin conformation data indicate a continuous global enhancement in topologically associating domains (TADs) boundary strength and increased chromatin accessibility at CTCF binding sites on these boundaries, identifying a new biomarker for brain aging. Additionally, our spatial transcriptome data underscore the variability in aging susceptibility even among the same cell type, emphasizing the importance of brain-region-level heterogeneity in unraveling the complexities of aging. Finally, we developed deep learning methods to accurately predict gene expression from multi-omic epigenetic data, underscoring the power of integrative computational approaches for advancing our understanding of aging biology. This study provides essential resources for examining the cellular, spatial, and regulatory genomic diversity within the aging mouse brain and supports the future development of a virtual brain aging model.

## RESULTS

### Integrative single-cell multi-omic dataset of aging mouse brain

We refined and employed snmC-seq3, an advanced single-nucleus methylome sequencing technique, to achieve base-resolution profiling of genome-wide 5-methylcytosine (5mC) across several brain regions in C57BL/6 male mice **(Figure 1A)**. These major regions, which include the anterior hippocampus (AHC), posterior hippocampus (PHC), frontal cortex (FC), amygdala (AMY), nucleus accumbens (NACB), PAG/PCG, entorhinal cortex (ENT), and caudate putamen (CP) **(Figure 1B)** (brian dissection details, see **Table S1)**, were meticulously dissected from male mice at 2, 9, and 18 months of age **(Figure 1A)**. Each region was sampled in two replicates, pooled from 3-7 different animals per replicate. Additionally, we employed snm3C-seq^26^, to assess the DNA methylome and chromatin conformation simultaneously in the same brain cell from the same regions of female mice **(Figure 1A)**. This approach integrates the 3D genome context across various cell types and ages, enriching our understanding of cellular changes over time. Single nuclei were captured using fluorescence-activated nuclei sorting (FANS), which enriched for neurons that were positively labeled with a NeuN antibody (NeuN+ neurons constituted 91.6% of snmC and 68.5% of snm3C data, with the remaining data being NeuN-neurons or non-neurons; Methods). Collectively, we obtained 132,551 DNA methylome profiles that passed quality control (QC). The snmC-seq data averaged 1.07±0.5 million final reads (mean ± s.d.) per profile **(Table S2)**. We also obtained 71,666 joint methylome and 3C profiles that passed QC, with each cell from this dataset having 1.9±0.5 million final reads (**Figure S1A)**. The 3C profiles revealed 442,000±156,000 cis-long-range (two anchors >2,500 bp apart) and 357,000±138,000 trans contacts per cell, respectively (**Figure S1B**, **Methods, Table S3**).

**Figure 1.**
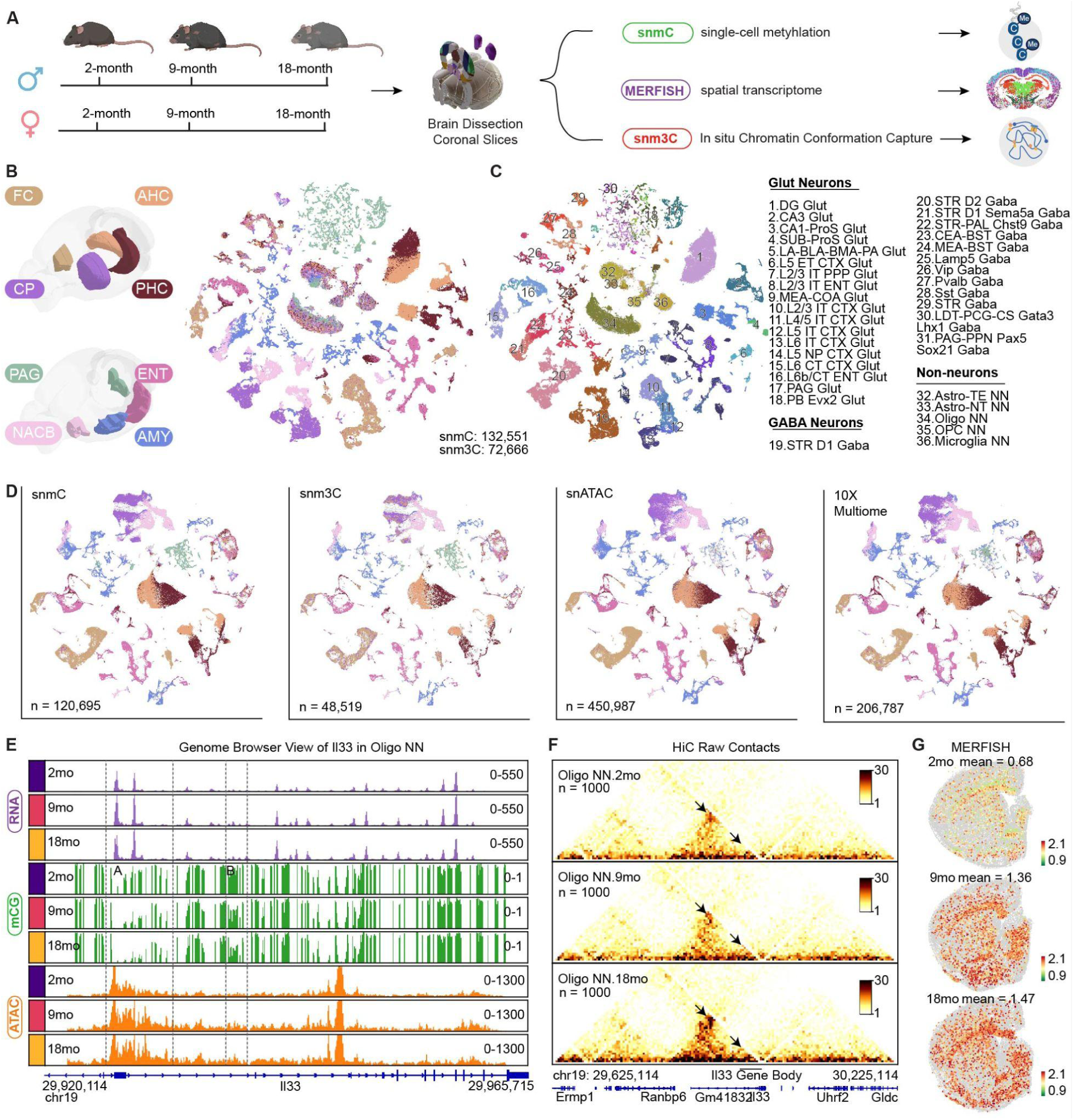
Single-cell multi-omic atlas reveals cell-type-specific aging signatures across brain regions. (**A**) Experimental workflow depicting age groups, tissue dissection methods, and molecular profiling approaches, including snmC-seq3, snm3C-seq, and MERFISH. (**B**) Left: Plot shows brain regions dissected and registered to the 3D Common Coordinate Framework (CCF)^101^; Right: Integration t-SNE of all cells from the snmC-seq3 and snm3C-seq, colored by brain regions. (**C**) Integration t-SNE of all cells from the snmC-seq3 and snm3C-seq, colored by cell types. (**D**) Integration t-SNE of all neurons from the snmC-seq (n = 120,695), snm3C-seq (n = 48,519), snATAC-seq (n = 450,987), and 10x Multiome (n = 206,787) datasets, colored by brain region. Light grey cells in the background represent cells from the other three modalities. snATAC and 10X multiome data were obtained from our companion study sharing the same brain samples. **(E-G)** The *Il33* gene exemplifies pseudo-bulk profiles of different modalities in “Oligo NN” across age groups, with a genome browser view showing RNA (purple), mCG (green), and ATAC (orange). Dash lines indicate genome regions (A and B) with notable changes in mCG and ATAC **(E);** heatmap of raw contacts around Il33 gene; arrows point to loop interactions with gene bodies **(F)**; spatial pattern of *Il33* gene expression on MERFISH coronal brain slice (**G**).

After QC and preprocessing, we clustered and integrated the snmC and snm3C datasets (Methods). Utilizing the annotations from the Brain Initiative Cell Census Network (BICCN)^27^, we identified 85 neuronal and 10 non-neuronal cell types **(Table S2 and S3)**. We focused on 36 major cell types, each represented by over 100 cells in every age group across both datasets, ensuring sufficient coverage for downstream analysis (**Figure 1C**). Consistent with previous observations^27,28^ from whole mouse brain atlases, global methylation levels varied substantially among cell types **(Figure S1C-S1E)**. Each cell type displayed mCH hypomethylation in its cell-type-specific marker genes **(Figure S2A)**. Within individual cell types, nuclei from different age groups show good alignment **(Figure S2B)**, achieving an alignment score^29^ of 0.98 ± 0.02 (Figure S2C, Methods).

To systematically identify alterations in gene expression and annotate the accessibility status of age-related differentially methylated regions (age-DMRs), we integrated our methylome data with our companion datasets. These included single-nucleus ATAC-seq (snATAC-seq) and 10x Multiome data generated from the same dissected samples used for snmC-seq3 and snm3C-seq analysis. We adopted our previously modified Seurat framework^27^ to integrate these datasets (Methods), which effectively matched neuronal spatial distribution **(Figure 1D)**, clusters **(Figure S2D),** and cell types **(Figure S2E)**. This integration generated cell-type-specific pseudo-bulk profiles spanning five molecular modalities: mCH/mCG fraction, chromatin conformation, chromatin accessibility, and gene expression.

To achieve higher spatial resolution and validate whether the observed epigenetic patterns correspond to transcriptomic changes, we employed MERFISH^30^ technology for *in situ* profiling of gene expressions in brain slices. We designed a 500-gene panel **(Table S4)** comprising 233 cell-type markers, selected based on cell type specificity and spatial diversity, along with 267 aging-related genes **(Methods)**. We profiled seven coronal sections from mice aged 2, 9, and 18 months. These slices include the brain regions included in our snmC and snm3C datasets. (**Figure S3A**). After quality control (**Figure S3B,** Methods), we obtained 895,296 MERFISH cells and assigned cell type annotations through integration with the BICCN whole mouse brain scRNA-seq^31^ dataset (**Figure S3C-S3F, Methods**, **Table S5**).

To illustrate the power of this comprehensive multi-omic aging dataset, we use the *Il33* gene, an inflammation gene known to be upregulated in aging oligodendrocytes (Oligo NN)^32^, as an example. Beyond confirming increased *Il33* RNA expression, we also demonstrate genome regions with a loss of methylation accompanied by gained chromatin accessibility during aging as putative cis-regulatory elements **(Figure 1E)**. The 3C heatmap revealed a higher frequency of contact, suggesting increased enhancer-promoter interactions that may regulate the expression of *Il33* **(Figure 1F)**. MERFISH spatial analysis further reveals the brain regions where *Il33* show upregulation, predominantly concentrated in white matter and subcortical areas (**Figure 1G**). Our study leverages single-cell methylation, single-cell chromatin conformation, and spatial transcriptome data to create a comprehensive genome atlas of the aging mouse brain spanning multiple modalities, brain regions, and cell types. This comprehensive resource has enabled numerous detailed analyses and led to various unique biological insights, which we will describe in the following sections.

### Age-related differential methylation signals in neurons and non-neurons

Our findings show that the genome-wide average methylation remains mostly stable across cell types **(Figure S4A and S4B)** and replicates **(Figure S4C and S4D)** in both sexes during aging, consistent with previous studies^33^. Specifically, we observed subtle global mCG DNA methylation changes when comparing 18-month-old to 2-month-old mice, with mCG fraction alterations of 0.008±0.005 in males and 0.006±0.006 in females across cell types (**Figure S4B**).

Despite relatively stable genome-wide average methylation levels, we identified age-related differentially methylated genes (DMGs) and differentially methylated regions (DMRs) at cell-type resolution. We identified 17,773 differentially CH-methylated genes (CH-DMGs) and 17,054 differentially CG-DMGs in young versus aged cells within cell types **(Methods)**. Gene body mCH and mCG had a strong negative association with mRNA expression in most cell types (**Figure S5A and S5C**, FDR < 0.01). For example, “L6 IT CTX Glut” showed a Pearson correlation coefficient (PCC) of −0.49 (FDR < 10^-30^) across DMGs, and “Oligo NN” cells showed a PCC of −0.58 (FDR < 10^-30^) (**Figure S5C)**. Notably, hypermethylated (aged > young) CG-DMGs were enriched in pathways related to ribosome complex biogenesis, ubiquitin-dependent protein catabolism, and mitochondrial ATP synthesis pathway, suggesting potential disrupted proteostasis and altered energy metabolism in aged brain cells **(Figure S5D-S5F)**. Interestingly, female striatum inhibitory neurons have the most abundant changes on X chromosome genes. These genes show decreased mCH methylation with aging **(Figure S5G**) and correlate with increased gene expression **(Figure S5H)**. For example, the *Dmd* gene encodes the inhibitory synapse scaffold protein dystrophin^34^ and is linked to intellectual disability^35,36^. This pattern aligns with previous observations in the female mouse hippocampus^37^.

DMRs are predictors of cis-regulatory elements, with hypomethylation of these regions typically indicating active regulatory status. We identified sex-shared methylation changes by combining reads from both sexes, then characterized age-DMRs in each major cell type across age groups (Methods). We identified more numerous and longer age-DMRs in non-neurons than neurons, independent of cell numbers (**Figure 2A**, **Figure S6A**). Among these, oligodendrocyte progenitor cells (OPCs) displayed the most abundant methylation changes. These findings align with previous observations^22,32^ for age-related transcriptome alterations, suggesting that non-neuronal cells are more substantially affected during aging. Specifically, we found 30,686±31,639 age-DMRs among non-neurons and 5,219±4,016 for neurons. These age-DMRs are highly cell-type-specific, indicating significant heterogeneity in aging among different cell types **(Figure 2B)**. Here, we define DMRs that progressively lose methylation with age as age-hypo DMRs (2mo > 9mo > 18mo), while those that continuously gain methylation are age-hyper DMRs (2mo < 9mo < 18mo). Neurons exhibited a higher proportion of intragenic age-DMRs compared to non-neuronal cells, and this pattern was consistent across age-hypo and age-hyper DMR categories **(Figure S6B)**. Notably, 88% of glutamatergic excitatory neurons exhibit a higher proportion of age-hypo DMRs **(Figure 2C)**. This may be partially explained by the significant decrease in *Dnmt3a* expression **(Figure S6C)**, a critical DNA methyltransferase for establishing new methylation patterns^38^. Both age-hypo and age-hyper DMRs show a strong negative correlation with chromatin accessibility status across age groups (FDR < 0.001) in most cell types **(Figure S6D and S6E)**, e.g., r = −0.8, FDR < 0.001 for “DG Glut” **(Figure S6E and S6F)**.

**Figure 2.**
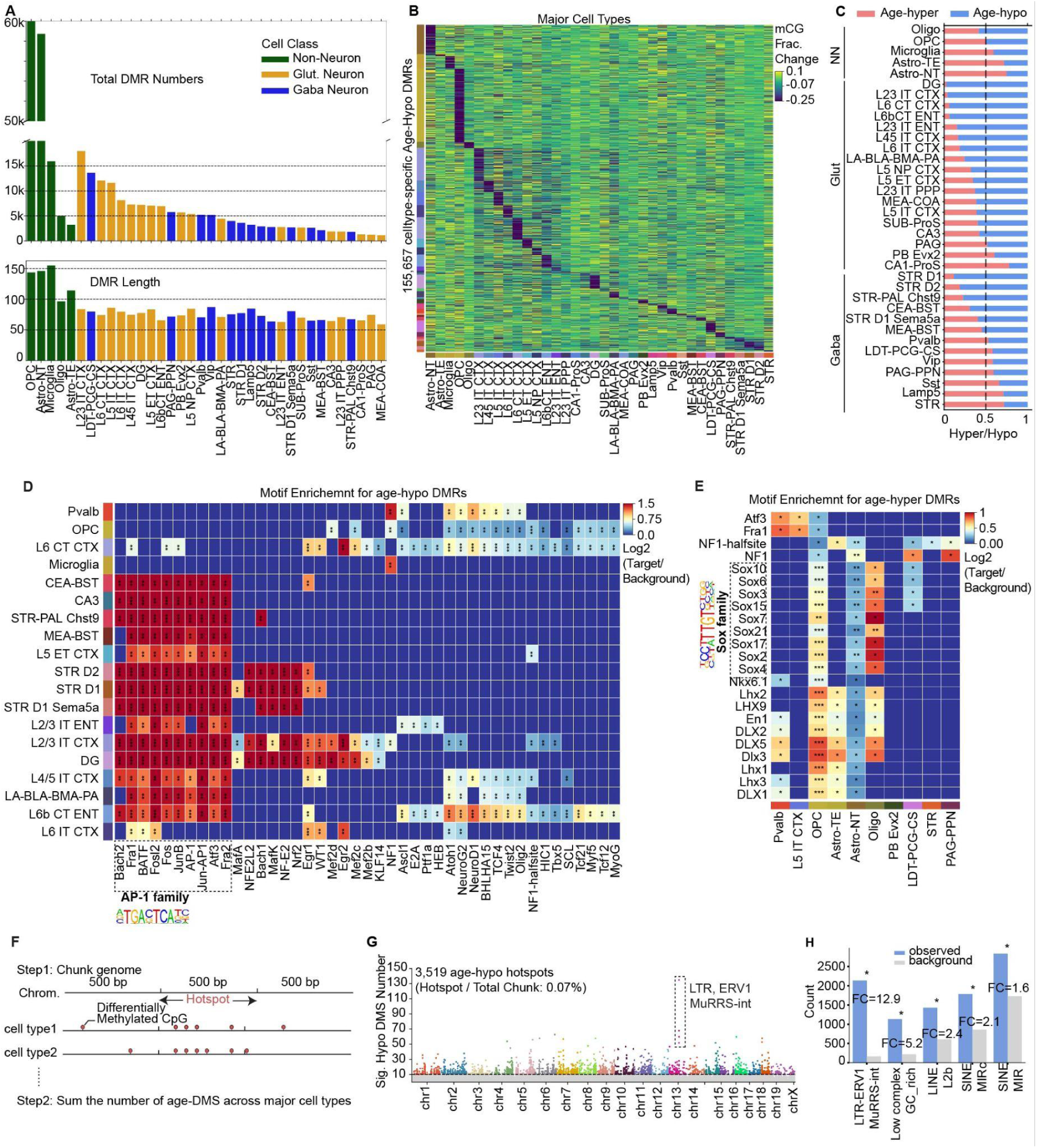
Age-related methylation alterations in neurons and non-neurons. **(A)** Bar plot shows the number of age-DMRs (top) and the average length (bottom) across major cell types. Bars are colored by cell class, including non-neuron (NN), glutamatergic excitatory neuron (Glut), and GABAergic inhibitory neuron (Gaba). **(B)** Heatmap depicts mCG fraction changes (ΔmCG: 18mo - 2mo) of age-DMRs (rows) across cell types (columns). The left color bar indicates the cell type source of the age-DMRs. **(C)** Bar plot shows the ratio of age-hypo and age-hyper DMRs across cell types, grouped by cell class (NN, Glut, and Gaba). **(D-E)** Heatmap shows the motif enrichment result using age-hypo **(D)** and age-hyper **(E)** DMRs. Color represents log2[% of Target Sequences with Motif/% of Background Sequences with Motif]. “*” indicate statistical significance (*P < 10^-5^, **P < 10^-30^, ***P < 10^-100^) **(F)** Schematic illustration of the genome aging hotspot identification strategy (Methods). **(G)** Scatter plot shows the number of age-hypo DMSs (y-axis) within 500 bp genomic chunks across chromosomes (x-axis). Each dot is a 500bp chunk, with hotspot chunks colored by chromosomes while other non-hotspot regions are shown in light grey. **(H)** Bar plot displays GAT (Methods) enrichment of TE subfamilies within age-hypo hotspot regions.

To explore the potential regulatory roles of DNA methylation changes, we performed sequence motif enrichment analysis **(Methods)** to identify transcription factors (TFs) that may bind at age-DMRs^39^. TF binding sites are mostly repressed if they gain DNA methylation^40^. We observed significant enrichment of AP-1 family motifs (for example, Fos, Jun) in neuronal age-hypo DMRs **(Figure 2D)**, consistent with observations in human cortical and hippocampal neurons^25,41^. Activation of AP-1 leads to chromatin opening, which disrupts neuron cell identity^42^ and has been linked to inflammaging in aged cells^41,43^. For age-hyper DMRs, motifs of the Sox transcription factor family, which is known for promoting self-renewal and differentiation^44^, are enriched in glial cells, including astrocytes, oligodendrocytes, and OPCs **(Figure 2E)**. Given that Sox10 is particularly crucial for OPC survival and development^45^, these results suggest a potential regulatory mechanism contributing to age-related decline in the OPC population observed in mouse **(Figure S6G)** and in human^41^, also their reduced differentiation potential^46^. In the companion paper (Amaral et al.), we discuss the dysregulation of transcription factors in different brain cell types during aging in greater detail.

### Retrotransposable elements emerge as methylation aging hotspots

We next investigated whether certain genomic regions are particularly enriched with aging signals. Methylation aging hotspots were identified as non-overlapping 500bp genomic regions containing more than 10 significant age-related differentially methylated sites (age-DMSs) across cell types **(Figure 2F, Figure S7A, Methods)**. The number of age-DMSs is not correlated with the CpG density **(Figure S7B)**. We identified 3,519 age-hypo and 969 age-hyper hotspots (**Figure 2G, Figure S7C**), representing regions with abundant age-associated methylation changes. Most hotspot regions are shared among cell types, whereas some represent ’cell-type unique hotspots’ characterized by dense signals (>80%) from a single cell type **(Figure S7D)**. We observed that 73.7% of age-hypo hotspots and 54.9% of age-hyper hotspots showed overlap with genes, which were subsequently termed “hotspot genes”. Interestingly, age-hypo hotspot genes show significant enrichment in GTPase-related processes, suggesting critical neuronal functions, including synaptic plasticity^47^ and autophagy^48^ **(Figure S7E)** might have been impacted. Surprisingly, when we checked the hotspots with the most hypomethylated signals, the top four hits were found in gene deserts and overlapped with retrotransposable elements on chromosome 13, specifically within the (Long Terminal Repeat) LTR class, ERV1 family, and MuRRS-int subfamily (LTR-ERV1-MuRRS-int) **(Figure 2G)**. We statistically checked the enrichment of TE subfamilies in age-hypo hotspot regions, highlighting a 13-fold enrichment in LTR-ERV1-MuRRS-int **(Figure 2H, Methods)**.

### TE methylation level distinguishes cell types and age groups

TEs are interspersed repetitive genetic sequences constituting a large portion of mammalian genomes, accounting for approximately 45% of human DNA and 40% of mouse DNA^49^. TE activity is tightly regulated through diverse epigenetic mechanisms; however, this control becomes compromised as cells age and TEs are reactivated^50^. This dysregulation of TEs can be detrimental through multiple mechanisms, including the induction of double-stranded DNA breaks^14^ and stimulation of inflammatory responses^17,51–55^.

DNA cytosine methylation represents a primary epigenetic modification that maintains transcriptional suppression of retrotransposons^14,50,56^. We initially focused on LINE regions, which are the most abundant retrotransposon class in humans. LINE1 elements are known to perform autonomous transposition in humans and contribute to age-related neurological disorders^57^. Our analysis revealed that intergenic and intragenic LINE regions function as effective features for clustering based on mCG methylation levels, demonstrating clear separation between cell types **(Figure 3A, Figure S8A, Methods)**. This suggests that even intergenic TE methylation states are critical functional features of cellular identity, independent of gene body methylation levels. Interestingly, we also observed pronounced segregation of age groups within cell types **(Figure 3A)**. The “DG Glut” exemplifies this age-related pattern most clearly, characterized by an increased ratio of hypomethylated LINE regions in aged neurons **(Figure 3B)** and the lowest alignment score^29^ between age groups **(Figure 3C)**. Surprisingly, when “DG Glut” cells were clustered separately, age-related separation appeared exclusively in cells from the anterior hippocampus (AHC) **(Figure S8B)**, revealing regional heterogeneity – a phenomenon explored in subsequent sections. Beyond “DG Glut”, other cell types with age group separations also exhibited an increased ratio of hypomethylated TE regions **(Figure S8C)**. These patterns were not unique to LINE elements, with similar methylation dynamics observed in LTR and Short Interspersed Element (SINE) regions **(Figure S8D and S8E)**.

**Figure 3.**
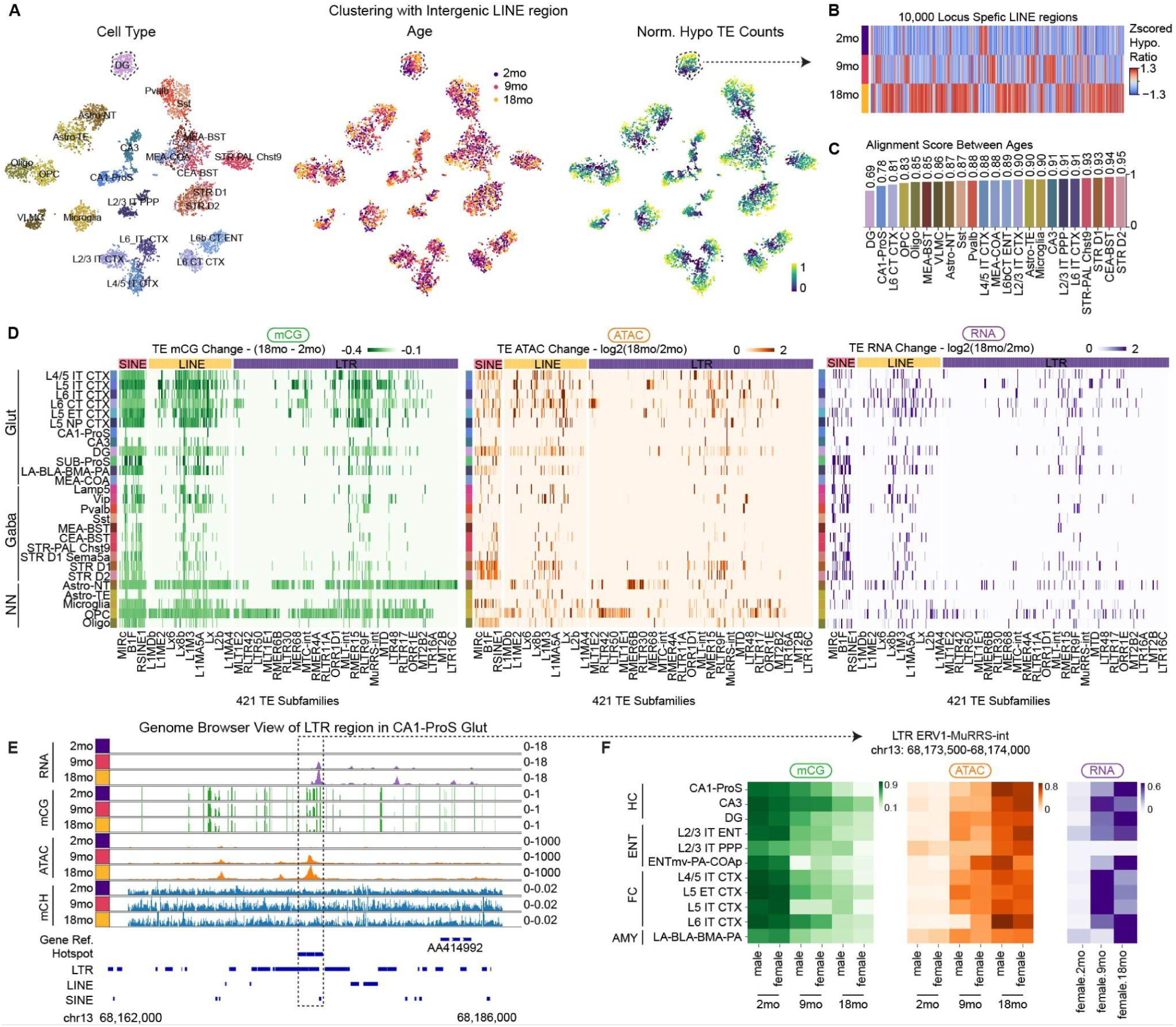
Genome-wide methylation profiling uncovers cell-type-specific retrotransposon activation. **(A)** t-SNE plot shows cell clustering based solely on Intergenic LINE regions. From left to right: t-SNE plots are colored by cell type, age, and the normalized count of hypo-methylated (mCG fraction < 0.75) LINE regions. **(B)** Heatmap exhibits the Z-scored TE hypomethylation (mCG fraction < 0.75) ratio among different age groups (rows) in “DG Glut”. Columns are the top 10,000 variant LINE regions. **(C)** Bar plot displays the alignment scores^29^ for each cell type across age groups, computed from the t-SNE dimensional reduction. Scores range from 0 (no age group overlap) to 1 (complete overlap). **(D)** Heatmap depicts age-related alterations in TE subfamilies (columns) across major cell types (rows). From left to right, the panels represent changes in mCG fraction (ΔmCG: 18mo - 2mo), chromatin accessibility (ATAC RPKM, Log2[18mo/2mo]), and gene expression (RNA RPKM, Log2[18mo/2mo]). **(E)** Genome browser visualization of pseudo-bulk profiles at the LTR hotspot region (chr13: 68,173,500-68,174,000) in “CA1-ProS Glut” across age groups. Tracks display RNA expression (purple), DNA methylation (mCG; green), chromatin accessibility (ATAC; orange), and mCH methylation (mCH; blue). Dashed lines highlight the LTR hotspot regions **(F)** Heatmaps display epigenetic and transcriptional profiles across cell types (rows). Each block shows, from left to right: DNA methylation (mCG), chromatin accessibility (ATAC), and RNA expression levels. Columns within each block represent different ages and sexes.

### Genome-wide TE Demethylation at cell type resolution

Previous studies^15,58^ typically examined TE methylation at the bulk level or using DNA array techniques that profile only targeted genomic regions. Our single-cell whole-genome bisulfite sequencing dataset enables investigation of genome-wide TE methylation patterns at cell type resolution. We conducted a systematic assessment of age-related methylation changes at specific genomic loci and found that 28.2±5.7% of age-hypo DMRs overlap with TE sequences. Our analysis revealed cell-type-specific methylation loss across 412 TE subfamilies, accompanied by correlated increases in chromatin accessibility and RNA expression **(Figure 3D)**. Excitatory neurons exhibit greater TE demethylation compared to inhibitory neurons, particularly among LTR regions (p < 0.05) **(Figure 3D**). A representative example is the LTR-ERV1 hotspot region on chromosome 13: 68,173,500-68,174,000. In “CA1-ProS Glut” neurons, this region exhibits age-associated demethylation concurrent with increased chromatin accessibility and RNA expression while maintaining stable mCH levels **(Figure 3E)**. Notably, this LTR region’s activation pattern is conserved across both sexes and is specific to excitatory neurons across multiple brain regions, including the hippocampus (HC), entorhinal cortex (ENT), frontal cortex (FC), and amygdala (AMY) **(Figure 3F, Figure S8F)**. These findings underscore that focusing solely on genes provides an incomplete picture of age-related changes, highlighting that the changes in the TE region are also crucial.

Next, we investigated the genomic locations where loss of TE methylation occurs. We found that regions undergoing TE methylation loss also lose H3K9me3^24^, a marker for heterochromatin **(Figure S9A and S9B),** suggesting that TE demethylation preferentially occurs in regions experiencing heterochromatin loss. For instance, L1MA5A in “CA1-ProS Glut” showed loss of methylation stemming from three different genome regions on chromosome 14, where they also exhibit decreased H3K9me3 levels **(Figure S9C)**. Moreover, the demethylation of TEs appears to be a continuous process **(Figure S9D)**, with the rate of methylation loss varying between the early (2 to 9 months) and late (9 to 18 months) aging stages **(Figure S9E)**. Cortex neurons, for instance, exhibit a faster rate of demethylation during early aging, whereas striatum (STR) inhibitory neurons display more significant changes during later stages. This suggests that TE methylation drifting rates are not uniform across different life stages. Next, we investigated the potential deleterious consequences of TE activation. Previous studies have demonstrated that TE depression leads to higher levels of cytoplasmic DNA^17^ or retrovirus-like particles^55^, triggering inflammation through cGAS-STING pathways. Our analysis revealed significant upregulation of genes involved in innate immune pathways across multiple cell types **(Figure S9F)**. Most notably, microglia, the brain’s primary immune cells, exhibited significant upregulation of key innate immune response genes, including *Nfkb1, Stat1,* and *Il18*^59–61^ **(Figure S9G)**.

### Increase in compartment strength during aging

Our annotated multi-omic aging datasets have enabled us to comprehensively investigate age-related dynamics of the 3D genome at cell-type resolution. We focused on systematic assessments of variability in key 3D genome features, including chromatin compartments, topologically associated domains (TADs), and loops. Our analysis reveals slight increases in short-to-long-range contact ratios and cis-to-trans contact ratios when comparing 2-month and 18-month-old samples, consistent with observations for human hippocampal neurons^41^ **(Figure S10A-S10C)**.

We expanded our analysis to the chromatin compartment, a crucial component of genome topology that organizes genomic regions spanning tens to hundreds of megabases^62^. The genome is organized into two primary compartments: A (active chromatin) and B (silent chromatin), reflecting distinct functional states^62^. Our study identified a total of 897 cell-type-specific age-related differential compartments (age-DCs), with 528 (58.9%) exhibiting switches between A-and-B compartments (**Figure S11A and S11B, Methods)**. These findings suggest substantial chromatin reorganization during aging, potentially impacting gene expression. For instance, chromatin rearrangements were observed on chromosome 2:168.3-173.8 Mb in oligodendrocytes. This region displays a switch from A to B compartments during aging, accompanied by reduced chromatin interaction frequencies **(Figure S11C)**. This reorganization correlated with significant downregulation of *Bcas1* **(Figure S11D**), a gene essential for early myelinating oligodendrocytes^63^. Notably, most excitatory neurons exhibited a higher ratio of B-to-A compartment switches **(Figure S11E)**, consistent with prior observations of heterochromatin loss in excitatory neurons^24^. We quantified compartment strength using the formula (AA + BB) / (AB + BA) and observed a consistent increase across cell types **(Figure S11F-S11H, Methods)**, in line with recent findings in cerebellar granular cells^20^. Specifically, excitatory neurons show an increase in B-B compartment interactions, while inhibitory neurons demonstrate more A-A compartment contacts **(Figure S11H)**. These results underscore both similarities and differences in compartment alterations across different cell types.

### Increase in TAD boundary probabilities and formation of smaller TADs

TADs are highly self-interacting chromosomal regions bounded by CTCF protein binding sites, which create insulated neighborhoods essential for gene regulation^64^. Together with cohesin, CTCF establishes chromatin loops that define these domain boundaries and regulate enhancer-promoter interactions^65^. Using contact matrices imputed at 25 kb resolution **(Methods)**, we identified TADs in single cells **(Methods)** and discovered a significant increase in the number of domains with a reduction in their average size during aging across different cell types (FDR < 10^-10^) **(Figure 4A)**. To explore the reason for the rise of smaller TADs, we analyzed age-related differential TAD boundaries (age-DBs) using data on boundary probability (Methods). Boundary probability means the proportion of cells where a genomic bin serves as a domain boundary. We identified a total of 2,469 cell-type-specific age-DBs showing significant variability (FDR < 1 × 10^-3^; Methods). Notably, 80.8% (1,994/2,469) of age-DBs were found in excitatory neurons, suggesting their particular susceptibility to TAD structural changes **(Figure 4B)**. Remarkably, 95.6% of these age-DBs exhibited increased boundary probabilities along with decreased insulation scores at the pseudo-bulk level **(Figure 4C)**, a trend that progresses continuously with aging **(Figure S12A)**.

**Figure 4.**
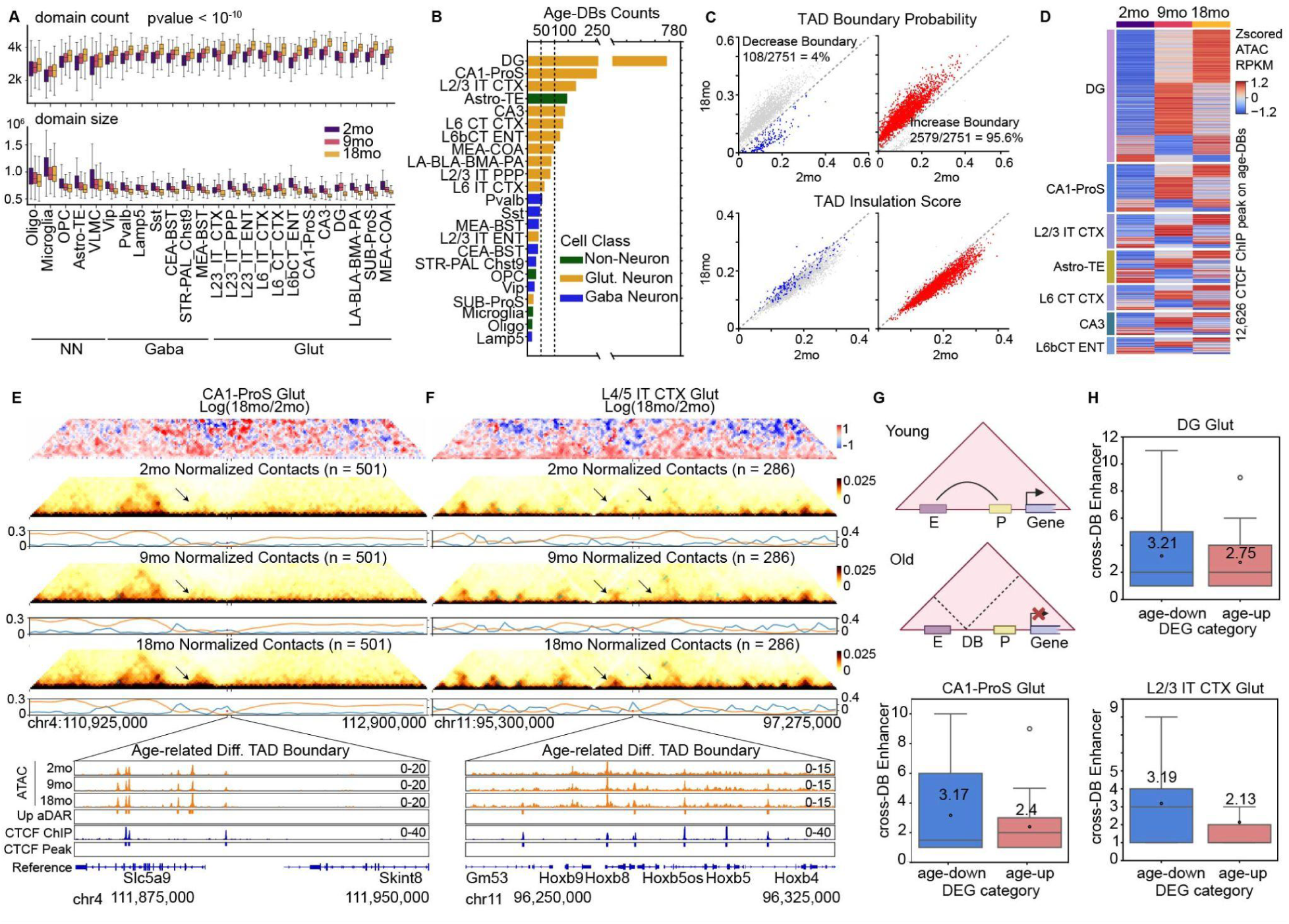
Age-related global increase in boundary probabilities associated with CTCF gaining accessibility. **(A)** Boxplots show the number (top) and size (bottom) of TADs across cell types in different age groups. Only cell types with significant increases in domain counts during aging (FDR < 1 × 10⁻¹⁰) are plotted. **(B)** Bar plot shows the number of age-related differential boundaries (age-DBs) across major cell types. Bars are colored by the cell class. **(C)** Scatterplot displays age-DBs comparing 2-month and 18-month timepoints. Boundary probability (top row) and insulation scores (bottom row) are plotted on the x-axis (2 months) and y-axis (18 months). Each point represents a single age-DB, with red and blue indicating increased and decreased boundary probabilities during aging, respectively. **(D)** Heatmap shows z-score ATAC RPKM at CTCF ChIP-seq peaks within age-DBs. The analysis compares chromatin accessibility patterns across cell types (rows) and age groups (columns). **(E)** Top, heatmaps show contact change (log2[18mo/2mo]) and normalized contact maps at 2, 9, and 18 months around the age-DB (chromosome 4: 111,875,000-111,950,000, extended 25,000 bp upstream and downstream) in “CA1-ProS Glut”. Orange and blue lines represent insulation score and boundary probability, respectively. Arrows indicate emergence of smaller TAD structure during aging. Bottom: genome browser view shows pseudo-bulk ATAC (orange) at the age-DB across age groups and CTCF ChIP-seq tracks. **(F)** Similar organization as in **(E)**, showing chromatin contact frequencies in “L4/5 IT CTX Glut” around the age-DB on chromosome 11: 96,250,000–96,325,000. **(G)** Schematic illustrating reduced promoter-enhancer interactions resulting from increased TAD boundary probability. **(H)** Boxplot displays the number of putative enhancers that cross age-DB (y-axis) for boundary proximal-genes located in the domain, with colors indicating expression changes (red: upregulated; blue: downregulated).

### Increase in chromatin accessibility on boundary CTCF binding sites

Next, we asked about the molecular mechanisms behind the observed global increase in TAD boundary probabilities. Given that CTCF is a crucial boundary insulator, we examined its potential contribution to this phenomenon. We first analyzed changes in chromatin accessibility at CTCF binding sites, where a previous observation showed that intra-TAD contacts correlate with CTCF accessibility during aging^41^. We utilized ENCODE^66^ mouse forebrain CTCF ChIP-seq data (Methods), which surprisingly revealed increased chromatin accessibility at 7,480 of 12,626 CTCF peaks on the age-DBs **(Figure 4D)**. We further investigated if DNA methylation might affect CTCF binding during aging. Previous in vitro fluorescence assays demonstrated that C2 methylation reduces CTCF binding affinity approximately 23-fold, while C12 methylation has no significant effect^67^. Additionally, single-molecule footprint analysis with ChIP-seq data indicated that C4 CpG also affects CTCF binding affinity^68,69^. Conversely, our analysis revealed a slight increase in average methylation levels at the C2 position of CTCF ChIP peaks, consistent with findings in human hippocampal cells^41^ **(Figure S12B)**. This minor change in methylation is likely attributable to the inherently low methylation status of cytosine sites within CTCF binding sites in young animals (mean: 0.036, median: 0) **(Figure S12C)**^70^. Next, we performed motif enrichment analysis of age-hypo DMRs and differentially accessible regions with increased accessibility (age-up DARs). Both analyses revealed significant enrichment of AP-1 (for example, Fos, Jun) family transcription factors **(Figure S12D and S12E)**. Significant CTCF enrichment was only observed in age-up DARs across cell types, mostly excitatory neurons **(Figure S12E and S12F)**. To illustrate the emergence of smaller TADs and changes in chromatin accessibility at age-DBs, we examined specific examples in “CA1-ProS Glut” and “L4/5 IT CTX Glut” cell types. In both examples, CTCF ChIP-seq peaks at age-DBs exhibited increased chromatin accessibility concurrent with elevated boundary probabilities and the formation of smaller TAD structures in the aged group **(Figure 4E and 4F)**.

We then investigated the potential transcriptional consequences of these changes. Previous studies have demonstrated that disrupting CTCF-associated TAD boundary elements leads to de novo enhancer-promoter interactions and gene misexpression^71^. Here, we hypothesized that increases in TAD boundary probabilities might lead to loss of enhancer-promoter interactions and subsequent transcriptional downregulation **(Figure 4G)**. To test this hypothesis, we first identified putative enhancers and their target genes using the activity-by-contact (ABC) model. We employed cell-type-specific ATAC peaks (from our companion study) and hypo-DMRs^27^, respectively as candidate cis-regulatory elements (cCREs), and used chromosome conformation data for linkage **(Methods)**. When examining genes proximal to age-DBs (boundary-proximal genes) **(Methods)**, we found that downregulated genes consistently exhibited a higher frequency of cross-boundary enhancer interactions in the young group, suggesting that age-related boundary strengthening may contribute to transcriptional decline by disrupting essential enhancer-promoter communications **(Figure 4H)**. Interestingly, these boundary-proximal genes are enriched in pathways regulating amyloid-beta formation **(Figure S12H)**. Collectively, these results highlight age-related TAD boundary strengthening as a novel molecular hallmark of aging. This boundary strengthening might be caused by CTCF peaks gaining chromatin accessibility and potentially impacts the expression of genes involved in Alzheimer’s disease pathogenesis.

### Regional variability of aging effects at cell-type resolution

The mammalian brain is composed of tens of millions of cells organized into intricate anatomical structures, each with unique cellular compositions and neural connectivity patterns. Previous studies have shown that cells of the same type exhibit substantial DNA methylation heterogeneity across different brain regions^27,28,72^. However, our understanding of how the same cell type ages across different brain regions remains poorly understood. To address this gap, we leveraged our single-cell, multi-region aging dataset to conduct a differential aging analysis, stratifying cells by cell type, region, and age. Our findings revealed notable regional variability in aging-related changes at the cell-type level. Specifically, we systematically quantified the number of age-DMRs **(Figure S13A)**, age-related differentially expressed genes (age-DEGs) **(Figure S13B)**, CG-DMGs **(Figure S14A)**, CH-DMGs **(Figure S14B)**, age-DBs **(Figure S15A)**, and age-related differential loops (age-DLs) **(Figure S15B)**.

To further illustrate regional heterogeneity in aging-related changes, we focused on “DG Glut”—granule cells located in the dentate gyrus that process sensory information critical for memory formation^73,74^, and also play a role in depression^75,76^. DG neurons exhibit distinct connectivity patterns along the anterior-to-posterior axis^77^. In our dissection experiments (Methods, Table S1), the anterior hippocampus corresponds to the dorsal hippocampus, while the posterior hippocampus includes the ventral regions plus part of the dorsal hippocampus. The dorsal hippocampus mediates cognitive functions, whereas the ventral regions are involved in regulating stress, emotion, and affect^78^. In our analysis, clustering of “DG Glut” cells by LINE regions revealed age-related separations that were exclusive to the anterior hippocampus (AHC), not the posterior hippocampus (PHC), suggesting a stronger influence of LINE elements on anterior DG granule cells **(Figure S8B)**. Further differential analysis indicated that age-associated alterations were more pronounced in AHC, with a higher abundance of age-DMRs, age-DEGs, age-DMGs, age-DBs, and age-DLsc **(Figure 5A)**.

**Figure 5.**
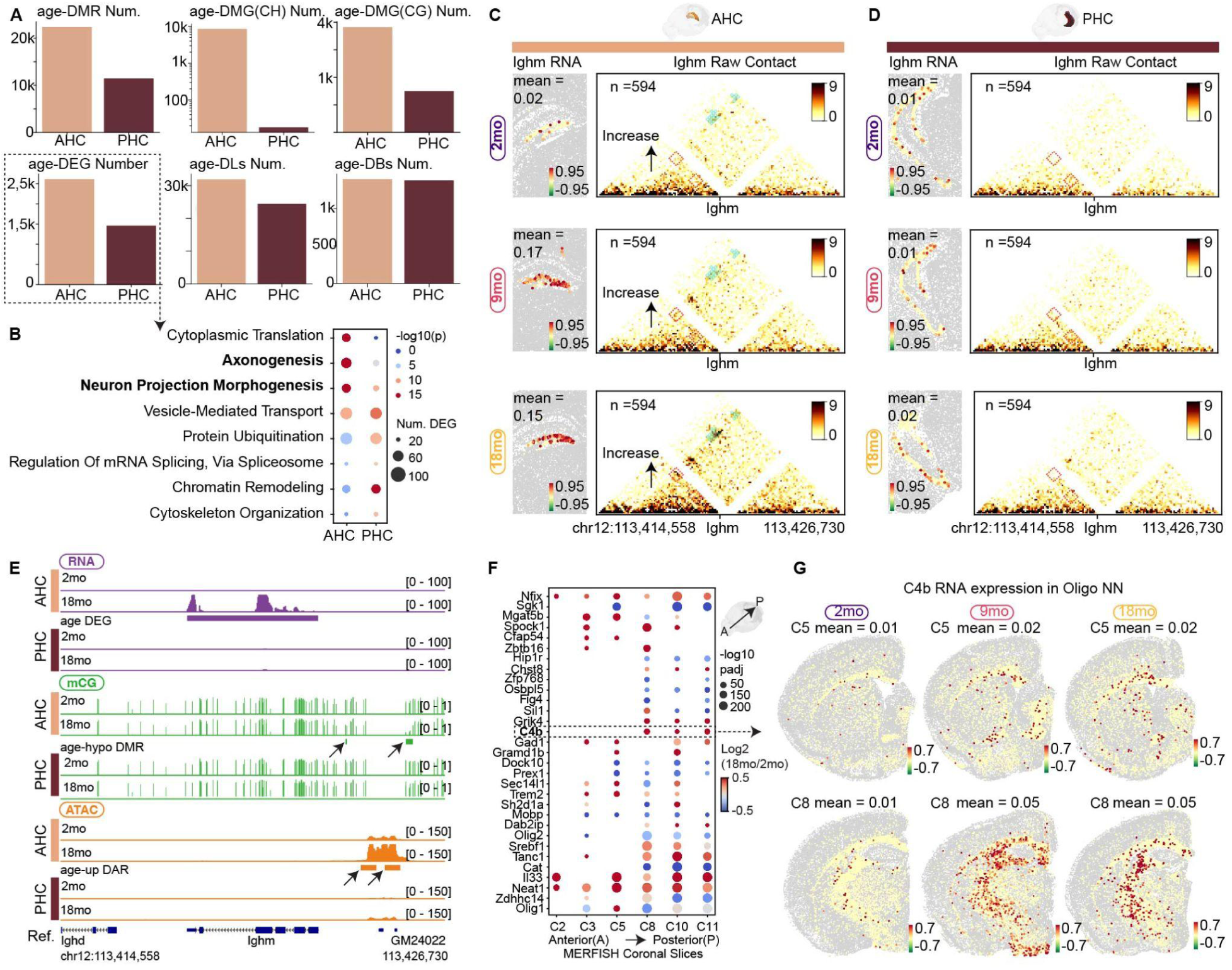
Spatial heterogeneity of aging signatures within identical cell types. **(A)** Barplots illustrate the quantification of age-associated changes of “DG Glut” in anterior (AHC) and posterior (PHC) hippocampus, showing the total number of age-DMRs, age-DMGs (using mCH or mCG), age-DEGs, age-DLs, and age-DBs. **(B)** Heatmap shows the GO pathway enrichment for age-DEGs in “DG Glut” neurons from AHC and PHC. Dot size represents the number of genes per pathway, and color intensity indicates statistical significance (−log10[P]). **(C)** *Ighm* expression in “DG Glut” on MERFISH slice C10 (anterior hippocampus): Top, plot shows AHC dissected and registered to the 3D CCF ^101^. Left column, spatial pattern of Ighm RNA expression across age groups (rows). Right column, heatmap of raw HiC contacts around the Ighm gene across age groups (rows). Arrows indicate contact strips spanning the Ighm gene body. **(D)** Similar to **(C),** corresponding information for “DG Glut” on MERFISH slice C11 (posterior hippocampus). **(E)** Genome browser view of RNA expression (purple), mCG fraction (green), and ATAC (orange) information around the *Ighm* locus across age groups in “DG Glut” neurons from AHC and PHC (rows). Arrows indicate age-DMRs and age-DARs. **(F)** Dotplot displays age-related transcriptional changes (rows) in “Oigo NN” across the anterior-to-posterior axis (left to right columns). Each dot represents an age-DEG on a specific MERFISH brain slice (columns), with size indicating statistical significance (−log10[P]) and color intensity showing the age-related transcriptional changes (log2[18mo/2mo]). **(G)** Spatial expression patterns of *C4b* in “Oligo NN” across age groups on MERFISH coronal slices C5 and C8.

Gene ontology enrichment analysis of AHC “DG Glut” age-DEGs identified in our companion study (Amaral et al.) showed significant enrichment in axonogenesis and neuronal projection pathways **(Figure 5B)**, suggesting aging may preferentially affect neuronal projections in anterior “DG Glut” neurons. We also compared gene expression of “DG Glut” neurons in AHC and PHC using our annotated MERFISH dataset **(Figure S3)**. Notably, the immunoglobulin heavy chain gene *Ighm* was upregulated in anterior hippocampal DG Glut neurons **(Figure 5C)**, but not in the posterior regions **(Figure 5D)**. This increased expression correlated with enhanced chromatin interactions observed solely in AHC **(Figure 5C and 5D)**, where upstream regulatory regions also showed DNA hypomethylation and increased chromatin accessibility **(Figure 5E)**. Age-related *Ighm* upregulation has also been observed in cortical excitatory neurons showing specific spatial patterns^32^. This upregulation of *Ighm* in anterior DG neurons may indicate heightened inflammatory stress in this neuronal population during aging.

Unlike neurons with their distinct anatomical organization, non-neuronal cell types are distributed throughout the brain^27,31,79^. Using anterior-to-posterior coronal slices from our MERFISH dataset **(Figure S3)**, we performed slice-specific differential expression analysis to identify age-DEGs (Methods). We first examined oligodendrocytes, the most prevalent non-neuronal cell type, which provide myelination to protect neuronal axons^80^. These cells exhibited marked regional variation in age-DEGs across brain regions. Notably, the *C4b* gene, a complement component and major schizophrenia risk factor^81^ and up-regulated in aged mice^82^ and neurodegeneration models^83^, showed selective upregulation in posterior brain slices **(Figure 5F and 5G)**. Spatial mapping showed that *C4b* upregulation was most pronounced in subcortical regions and white matter of the corpus callosum **(Figure 5G)**. Our analysis also revealed extensive regional heterogeneity in aging across other non-neuronal cells. For instance, the *Nlrp3* gene, a component of the inflammasome that detects cellular damage and initiates inflammation^84^, showed significant upregulation specifically in posterior microglias **(Figure S16A)**. Similarly, *Irf1*, implicated in cytokine interferon production^85^, was upregulated in astrocytes and OPCs exclusively in the posterior slices **(Figure S16B-D)**. These findings clearly demonstrate that both neuronal and non-neuronal cells display region-specific aging signatures even within the same cell type.

### Deep learning model predicts age-related gene expression changes using epigenetic data

Using our comprehensive multi-omic dataset, we developed machine learning frameworks to integrate diverse epigenetic modalities and uncover potential nonlinear relationships of gene expression changes. In our EpiAgingTransformer (EAT) model, we incorporate three epigenetic modalities—DNA methylation, chromatin accessibility, and chromatin conformation—each contributing distinct epigenetic features, for a total of seven features (𝑋). Specifically, DNA methylation includes gene body age-DMRs, as well as gene body average mCG and mCH levels. Chromatin accessibility is measured by ATAC-seq peaks within gene bodies. Chromatin conformation encompasses chromatin loops and putative enhancers identified through the ABC model using peak or hypo-DMR as cCREs **(Figure 6A and 6B, Methods)**. Collectively, these seven features provide a comprehensive representation of the epigenetic landscape as inputs to the EAT model. The model outputs gene expression changes during aging (𝑌) as −1 for age-downregulated genes, 0 for non-differentially expressed genes (non-DEGs), and 1 for age-upregulated genes **(Figure 6A, Methods)**. Our objective is to assess whether these transcriptome changes (𝑌) can be accurately predicted using the specified seven epigenetic features (𝑋).

**Figure 6.**
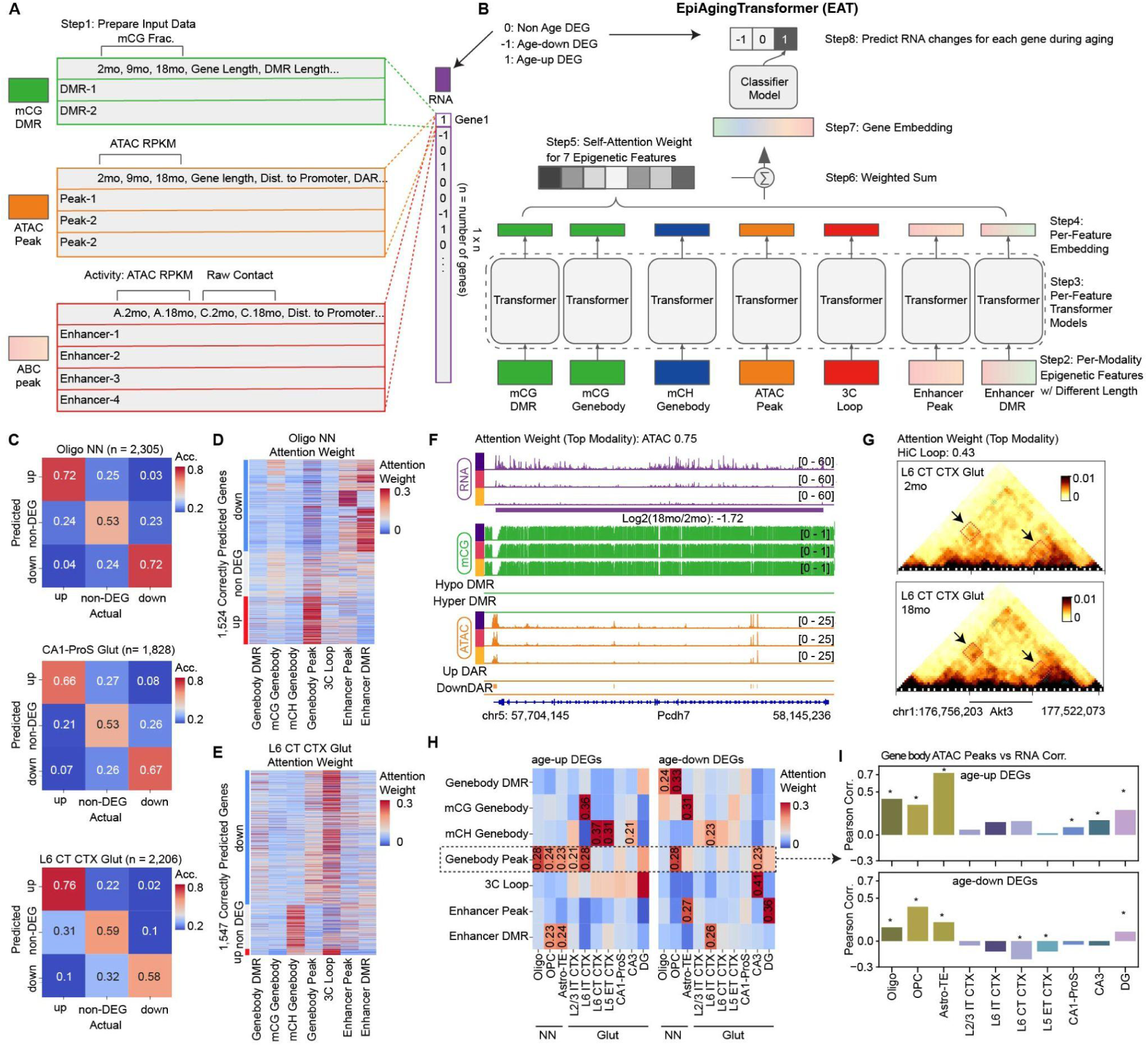
Deep learning model reliably predicts aging effects on gene expression using multi-modal epigenetic data. **(A-B)** Schematic representation of the input data **(A)** and the deep learning training workflow **(B)**. **(C)** Heatmap illustrates prediction accuracy across different cell types. The columns and rows represent actual and predicted categories, respectively. Color intensity and text indicate the accuracy for each gene category combination. **(D-E)** Heatmaps display attention weights from each epigenetic feature (columns) across all correctly predicted genes (rows) in “Oligo NN” **(D)** and “L6 CT CTX Glut” **(E)**. The sum of each row is 1. **(F)** Pseudo-bulk profiles of *Pdch7* in “Oligo NN” across age groups, alongside the genome browser view showing RNA (purple), mCG (green), and ATAC (orange). **(G)** Heatmap of raw contacts around the *Akt3* gene in “L6 CT CTX Glut.” Arrows point to differential loop interactions with gene bodies. **(H)** Heatmap depicts the average attention weight from each epigenetic feature (rows) across cell types (columns). Numbers on the heatmap indicate the attention weights if greater than 0.2. **(I)** Pearson correlation coefficients between changes in chromatin accessibility (log2[18mo/2mo]) and gene expression (log2[18mo/2mo]) across cell types. Top: age-upregulated DEGs; down: age-downregulated DEGs. “*” indicates statistically significant correlation (P < 0.05). Bars are colored by cell type.

The EAT model comprises seven transformer modules, each tailored to handle one epigenomic feature in data samples, which is crucial given the diverse lengths of regulatory elements across genes. This model architecture also includes a self-attention aggregator^86^ to aggregate the information from multiple features, which enhances the interpretability by assigning attention weights to seven epigenetic features. Last, we include a multiple-layer perceptron (MLP) classifier to predict the age-related differential expression **(Figure 6B, Methods)**. We evaluated the model performance using 5-fold cross-validation, training on four folds and predicting on the held-out fold to avoid data leakage **(Methods)**. Our EAT model effectively predicts age-related gene expression alterations using epigenetic information. After training the model separately across 10 cell types, it achieved an average accuracy (correctly predicted/total age-DEGs) of 74.5±8.5% for age-DEGs **(Figure 6C, Figure S17A)**. However, when the model was trained using data from single modalities, a notable decrease in accuracy was observed **(Figure S17B)**. For example, the “L6 CT CTX Glut” accuracy fell by 4.5-6.7% using only one modality **(Figure S17B)**.

In parallel, we employed a Random Forest (XGBoost) model **(Figure S18A, Methods)**, which also demonstrated comparable predictive power with an accuracy of 66.7±4.9% **(Figure S18B)**, further supporting that epigenetic features are reliable predictors of transcriptomic changes. Compared to a baseline linear model, specifically the logistic regression model for classification, which achieved an accuracy of 61.4±5.7% **(Figure S18C)**, the EAT and XGBoost models exhibit higher accuracies by 1.21x and 1.08x, respectively. These results suggest that the expression level of different genes is influenced by varying epigenetic features and that regulatory relationships are not uniformly linear.

### Model interpretation reveals biological insights into gene regulation during aging

To understand how different epigenetic modalities and features contribute to gene expression changes, we analyzed the attention weights from the self-attention aggregator. These weights provide insight into each feature’s relative contribution to our predictions. Our analysis indicates that genes display distinct patterns of feature-specific weights, suggesting varied regulatory mechanisms across genes **(Figure 6D and 6E, Figure S19A)**. For example, *Pcdh7*, a gene essential for cell-cell recognition and synaptic function and implicated in brain disorders^87^, shows significant age-related downregulation in “Oligo NN”. The model assigns the highest attention weight of 0.75 to its ATAC peak, with genome browser views confirming that changes in *Pcdh7* expression are predominantly driven by alterations in chromatin accessibility **(Figure 6F)**. Another example is the gene *Ppfibp2*, which is crucial for cell adhesion^88^, exhibits age-related upregulation in “Oligo NN”. This upregulation is primarily influenced by gene body age-DMRs, highlighted by a significant attention weight of 0.54 and the presence of a higher abundance of age-hypo DMRs **(Figure S19B)**. Another compelling example is *Akt3*, a kinase critical for neuronal survival^89^ associated with schizophrenia and bipolar disorder^90,91^. In “L6 CT CTX Glut”, the model assigns a substantial weight of 0.43 to the 3C loop feature, reflecting the significantly increased contact interactions associated with this gene **(Figure 6G)**. These examples exemplify how the self-attention module in the EAT model effectively quantifies the relative contributions of different epigenetic features to gene expression changes, validating the model’s capability in dissecting complex epigenetic regulation.

To systematically characterize these cell-type-specific regulations, we calculated the average attention weight for each epigenetic feature across gene categories **(Figure 6H)**. Notably, mCH gene body weights were predominantly allocated to neuronal cells, with neurons exhibiting higher average attention weights (17.6 ± 11.1%) compared to non-neuronal cells (7.6 ± 3.6%) **(Figure 6H)**. This observation aligns with previous studies indicating that non-CpG cytosines are abundantly methylated uniquely in mouse and human neurons during development^13,92^, and we extend this finding to the context of aging. Further, our analysis uncovered gene body ATAC peaks that were more strongly associated with age-related gene upregulation (average attention weight: 20.2 ± 5.9%) than with downregulation (16.8 ± 5.3%). This finding can be further validated that age-up DEGs demonstrate significantly (p < 0.01) higher correlation between ATAC peaks and RNA signals than downregulated DEGs **(Figure 6I)**. Specifically, up-regulated genes exhibit an average Pearson correlation coefficient of 0.24±0.2, whereas down-regulated genes show a substantially lower correlation of 0.03±0.21 **(Figure 6I)**. Conversely, changes in enhancers had a greater impact on gene downregulation during aging (15.2 ± 7.5%) compared to upregulation (11.9 ± 6.5%) **(Figure 6H)**. Collectively, our study uniquely establishes a comprehensive and interpretable machine learning framework for understanding age-related transcriptomic changes through the integration of multiple epigenetic modalities and features.

## DISCUSSION

Aging in the brain presents a significant health challenge, with epigenetic dysregulation emerging as a pivotal mechanism. However, the functional significance of methylation clock sites remains unclear without comprehensive genome-wide analysis. Prior transcriptomic data have revealed widespread age-related changes, typically pointing toward increased inflammation^93,94^ and decreased neurogenesis^22^—increased inflammation and reduced neurogenesis, yet the epigenetic drivers behind these shifts are not fully understood.

Here, our single-cell multi-omic data reveal fundamental aspects of epigenetic aging in the mouse brain, including heightened vulnerability of non-neuronal cells to age-related methylation changes, cell-type-specific transposable element demethylation, and a pronounced increase in the number of smaller topologically associating domains (TADs)—particularly in excitatory neurons. We also highlight the spatial heterogeneity of aging processes across different brain regions. Taken together, these findings underscore the cell-type- and region-specific organization of the aging mouse brain, which we discuss in greater detail below.

Consistent with recent findings on differentially expressed genes in aging brains^22,32^, we observed that non-neuronal cells exhibit more pronounced age-related methylation changes than neurons. Both transcriptomic and methylomic data point to glial cells as especially susceptible to age-related alterations. Motif analysis of age-DMRs revealed robust AP-1 family activation in neurons, a phenomenon also reported in hematopoietic stem cells^95^, kidney and liver^43^, suggesting a conserved role for AP-1 in aging. Prior studies^42^ suggest that AP-1-linked chromatin opening disrupts cell identity TF binding. Notably, we also detected increased methylation at Sox transcription factor binding motifs in OPCs. Given Sox’s critical role in OPC survival and terminal differentiation^45^, alterations in Sox motif methylation could compromise oligodendrocyte maturation and subsequent axonal protection. Our findings suggest that targeting epigenetic regulation in OPCs at Sox motifs might preserve myelin integrity and mitigate neuronal degeneration in the aging brain.

DNA methylation represents a primary epigenetic modification that maintains transcriptional suppression of retrotransposons. Our single-cell genome-wide analysis demonstrates that TE methylation patterns can distinguish both cell type and aging status, suggesting TEs can serve as functional elements and have implications in aging. Furthermore, we identified correlations among retrotransposon hypomethylation, increased chromatin accessibility, and elevated RNA levels, validating the age-related TE activation. Previous studies also showed that chromatin derepression can be exacerbated by L1 RNA overexpression^18^, which enables further TE activation and triggers innate immune response and inflammation^16,17^. Based on these findings, we propose that TE activation may initiate a self-reinforcing cycle, though the initiating event remains unclear. Notably, clearance of LINE-1 cDNA has been demonstrated to reduce inflammation and DNA damage^16^ and extend mouse lifespan^17^. Our results can facilitate future interventions aimed at selectively silencing pathogenic TE activation patterns while preserving regulatory TEs vital for normal gene expression.

We also present the most comprehensive single-cell chromatin conformation dataset on the aging brain to date. Previous work^20^ observed increased chromatin compartmentalization during aging in mouse and human cerebellar granule cells; our study extends this finding across multiple cell types, suggesting a general mechanism. Our findings also uncover a unique biomarker of aging, characterized by a significant increase in the number of smaller TADs across most cell types, particularly in excitatory neurons. Moreover, a substantial fraction of TAD boundaries exhibit enhanced boundary probabilities. This increase correlates with increased chromatin accessibility at CTCF sites within these boundaries. Interestingly, the increase in CTCF chromatin accessibility and intra-TAD contacts is also observed in human hippocampus dentate gyrus (DG) granule cells^41^, suggesting a conserved mechanism. The strengthened TAD boundaries form more defined and isolated TADs, potentially interfering with promoter-enhancer interactions and explaining the downregulation of genes with cross-boundary enhancers.

Our multi-region analysis reveals considerable spatial heterogeneity in brain aging, underscoring the necessity to consider aging as a region- and cell-type-specific process. For instance, DG granule cells in the anterior hippocampus exhibit more pronounced changes compared to the posterior hippocampus, with anterior-specific upregulation of Ighm accompanied by promoter hypomethylation and enhanced chromatin accessibility. A previous study^96^ confirmed Ighm transcripts’ translation in neurons. These proteins may serve as cell adhesion molecules^97^ or activate the complement system, potentially contributing to brain immune responses^98,99^. Beyond neurons, non-neuronal cells also exhibit regional heterogeneity in aging. Previous spatial studies^32,100^ on aging have pointed out that white matter is particularly vulnerable to age-related changes. We demonstrated this spatial heterogeneity in greater detail, showing that C4b, a complement component and key risk factor for schizophrenia^81^ are only upregulated in the posterior white matter.

Our success in developing the EpiAgingTransformer (EAT) model demonstrates the power of integrative approaches for deciphering brain aging complexity. By successfully predicting age-related gene expression changes using multiple epigenetic modalities, EAT leverages self-attention mechanisms that enable direct interpretation of each epigenetic feature’s contribution. Analysis of these attention weights revealed unique biological insights: mCH modifications predominantly influence neuronal cells while having a minimal impact on non-neuronal populations. In addition, chromatin accessibility exhibits stronger effects on upregulated age-DEGs compared to downregulated ones. This multimodal deep learning framework provides a comprehensive view of how epigenetic factors orchestrate transcriptional changes throughout the aging process.

In conclusion, we present a comprehensive, rigorously validated single-cell multi-omic atlas of the aging mouse brain, illuminating the epigenetic brain aging landscape. Our findings reveal cell-type-specific transposon demethylation, increased TAD boundary strength, and notable spatial heterogeneity. Additionally, we developed a deep-learning model leveraging the multi-omic training sets to predict gene expression accurately. Translating these findings to humans may offer valuable avenues for targeted therapies and potentially constructing a “virtual” aging brain model.

## Limitations of the study

Our study faces limitations such as the difficulty in mapping multi-mappable transposable element reads to specific genome positions due to short-read sequencing and a technical inability to detect transposable element mRNAs lacking poly-A tails. Additionally, using different sequencing technologies for male and female samples complicates reliable sex-based comparisons in brain aging.

## Resource availability

### Lead contact

Further information and requests for resources and reagents should be directed to Joseph R. Ecker.

### Materials availability

This study did not generate new unique reagents.

### Data and code availability

The raw data from 2-month-old male mice have been deposited in NCBI’s Gene Expression Omnibus (GEO) database under accession numbers GSE132489 and GSE213262. Processed data are available under accession number GSE292803. All analysis code and processed data related to the results and methods sections are available on GitHub at https://github.com/rachelzeng98/aging-mouse-brain.git. The machine learning training code and processed input data are also available on GitHub at https://github.com/Michaelvll/aging-gene-prediction. Any additional information needed to reanalyze the data reported in this paper can be obtained from the lead contact upon request.

## Supporting information

Table S1. Brain Dissection Region Annotation

Table S2. Metadata for snmC-seq3

Table S3. Metadata for snm3C-seq

Table S4. MERFISH gene panel

Table S5. Metadata for MERFISH

Table S6. Major Cell Type Annotation Table

## Acknowledgements

This work was supported by the National Institutes of Health/National Institute on Aging (NIH/NIA) grant 5R01AG066018-05 to J.R.E and B.R. The Flow Cytometry Core Facility of the Salk Institute (RRID: SCR_014839) is supported by funding from NIH-NCI CCSG P30 CA014195, and Shared Instrumentation Grants S10-OD023689, and S10 OD034268. J.R.E. is an investigator of the Howard Hughes Medical Institute.

## Author contributions

J.R.E., Q.Z., B.R., and M.M.B. conceived the study. Q.Z., W.T., A.K., Z.W., Y.S., W.W., and W.D. analyzed the data. Q.Z., H.C., and J.R.E. uploaded the data. Q.Z., A.K., and Z.W. drafted the manuscript. J.R.E., B.R., M.M.B., H.L., and Y.R.L. edited the manuscript. J.R.E., Q.Z., M.M.B., and B.R. coordinated the research. J.R.E., M.M.B., J.O., N.D.J., R.G.C., J.R.N., J.A., M.K., C.V., and A.B. generated the snmC-seq3 and snm3C-seq data. M.M.B., W.O., C.T.B., A.A., S.C., C.C., J.W., S.C., J.R., J.L., A.B., J.A. generated the MERFISH data. B.R. and M.L.A. provided the snATAC-seq and 10X Multiome data. Q.Z. and H.C. contributed to the data archive. J.R.E., M.M.B. supervised the study.

## Declaration of interests

J.R.E. is a scientific adviser for Zymo Research Inc., Ionis Pharmaceuticals, and Guardant Health. B.R. is a cofounder and consultant for Arima Genomics Inc. and cofounder of Epigenome Technologies.

## Methods

### Mouse brain tissues

All experimental procedures using live animals were performed at the Salk Institute and were approved by the Animal Care and Use Committee (IACUC). The Salk Institute has approved Assurance from NIH Office of Protection from Research Risks (D16-00328). Male and female C57BL/6J mice were purchased from Jackson Laboratories at 7, 38 and 63 weeks of age and maintained in the Salk animal barrier facility on 12-h dark–light cycles with food ad libitum until brain extraction (housing conditions: temperature of 21–23 °C, relative humidity of 61–63%). For synchronizing the estrous cycle in females ahead of brain extraction, females were exposed to male bedding for three days such that they would be in di-estrous at the time of dissection. All animals were dissected between 1 and 5 pm to ensure similarity of circadian rhythm. Brain extraction, slicing, and dissection were performed in an ice-cold dissection buffer as previously described^28^. For both snmC-seq3 and snm3C-seq samples, we prepared coronal brain slices at 600 μm intervals spanning the entire brain, producing 18 slices total. Brain regions were dissected according to the Allen Brain Reference Atlas Common Coordinate Framework version 3 (CCFv3)^103^. Each region was collected from 2-7 animals and pooled to create 2-3 biological replicates for nuclei isolation. Sample size was determined empirically rather than through statistical methods. We established that two to three biological replicates for all single-cell epigenomic experiments would provide minimum reproducibility standards for this large-scale project. We did not implement blinding or randomization during tissue sample handling. All dissected regions underwent digital registration into CCFv3 using ITK-SNAP^104^ (v4.0.0) at 25 μm resolution.

### Nuclei isolation and Fluorescence Activated Nuclei Sorting (FANS)

Fluorescence-Activated Nuclei Sorting (FANS) using a BD Influx sorter in single-cell mode (1 drop single) to separate neuronal from non-neuronal populations at a ratio of 90:10 (NeuN+:NeuN-), following established protocols as previously described^28^. Nuclei were sorted into 384-well plates according to previously described methods^28^. The snm3C-seq protocol incorporated additional chromatin conformation capture steps during nuclei preparation to enable assessment of chromatin architecture. These steps utilized the Arima-3C BETA Kit (Arima Genomics), with comprehensive methodology detailed in the our previous publication^27^.

### Library preparation and Illumina sequencing

Both snmC-seq3 and snm3C-seq samples underwent identical library preparation procedures as detailed in the our previous publication^27^. We implemented automated workflows using Beckman Biomek i7 and Tecan Freedom Evo platforms to accommodate high-throughput processing requirements. Libraries were sequenced on an Illumina NovaSeq 6000 system, with each S4 flow cell processing 16 384-well plates using 150 bp paired-end sequencing parameters. The experimental workflow utilized several specialized software applications: BD Influx Software v1.2.0.142 for flow cytometry analysis, Freedom EVOware v2.7 for automated library preparation, Illumina MiSeq control software v3.1.0.13 and NovaSeq 6000 control software v1.6.0/RTA v3.4.4 for sequencing operations, and Olympus cellSens Dimension 1.8 for image acquisition and processing.

### Mapping and primary quality control

We performed snmC-seq3 and snm3C-seq data processing using the YAP pipeline (cemba-data package, v1.6.8). The mapping workflow began with demultiplexing FASTQ files to separate individual cells using cutadapt (v2.10)^105^, followed by read-level quality control assessment. Sequence alignment utilized one-pass mapping for snmC and two-pass mapping for snm3C with hisat3n (v2.2.1) and bowtie2 (v2.3)^106^. We then processed and quality-controlled BAM files using samtools (v1.9)^107^ and Picard (v3.0.0), generated methylome profiles with allcools (v1.0.8), and identified chromatin contacts exclusively for snm3C-seq samples. We have provided Snakemake^108^ pipeline files with comprehensive mapping details in the “Code availability” section. All sequence reads were aligned to the mouse mm10 reference genome. For gene and transcript annotation, we employed a customized version of the GENCODE vm22 GTF file.

For DNA methylome quality control, we applied several primary filtering criteria. Cells were required to have: (1) overall mCCC level < 0.05; (2) overall mCH level < 0.2; (3) overall mCG level > 0.5; (4) total final reads > 500,000 and < 10,000,000. The mCCC level serves as an upper boundary estimate for cell-level bisulfite non-conversion rate. We also calculated lambda DNA spike-in methylation levels to estimate non-conversion rates for each sample. Subsequent clustering-based quality control identified potential doublets and low-quality cells. For the snm3C-seq cells, we also required cis-long-range contacts (two anchors > 2500 bp apart) > 50,000.

### Analysis infrastructures

The scale of our dataset significantly exceeded previous bulk and single-cell studies examining methylation and chromatin conformation. To manage this complexity efficiently, we implemented a cell-by-feature tensor framework called “Methylome Cell DataSet” (MCDS) for methylome-based clustering and cross-modality integration. This approach concentrated on individual cells with aggregated genomic features such as kilobase chromosome bins, gene bodies, and transposon element regions. By aggregating single-cell profiles into pseudo-bulk levels through clustering and annotation, we achieved enhanced genome coverage while eliminating the need to repeatedly access hundreds of terabytes of single-cell files during subsequent analyses. We stored these large tensor datasets using the chunked and compressed Zarr format ^109^. Data analysis relied on ALLCools^28^, Xarray^110^, and dask^111^ packages. For efficient large-scale computation, we constructed pipelines using the Snakemake package ^108^ and established cloud environments with the SkyPilot package^112^. We updated the ALLCools package (v1.0.8) to perform methylation-based cellular and genomic analyses, and enhanced the scHiCluster^113^ package (v1.3.2) for chromatin conformation analysis. Comprehensive heatmaps presented were generated using the PyComplexHeatmap package (1.7.4)^114^. Complete tensor storage information and reproducibility details, including package versions, analysis notebooks, and pipeline files, are provided in the “Data and Code availability” section.

### Methylome-based Clustering

Following mapping, we stored single-cell DNA methylome profiles from both snmC-seq and snm3C-seq datasets in the “All Cytosine” (ALLC) format—a tab-separated table compressed and indexed using bgzip/tabix^115^. We then utilized the “generate-dataset” command in the ALLCools package to create a methylome cell-by-feature tensor dataset (MCDS). For clustering analysis, we employed non-overlapping 100Kb chromosome bins (chrom100k) of the mm10 genome as features. For both snmC and snm3C datasets, we used their mCH and mCG fractions of chrom100k for clustering. Detailed Jupyter notebooks documenting the functions and parameters for each analytical step are available in the “Data and Code availability” section. The majority of our analytical functions were derived from the allcools^28^, scanpy^116^, and scikit-learn^117^ packages.

Our clustering pipeline includes several key steps. We began with basic feature filtering based on coverage metrics and the ENCODE blacklist^118^. Next, we generated posterior chrom100k mCH and mCG fraction matrices, similar to a previous study^28^ and initially established in Smallwood et. al.^119^. We then calculated principal components (PCs) and created t-SNE and UMAP^120^ embeddings for visualization purposes. The t-SNE analysis was conducted using the openTSNE^121^ package, implementing procedures as previously described^122^. Finally, we performed consensus clustering by applying the Leiden algorithm^123^.

### Atlas-level data integration and cluster annotation

We established a highly efficient framework based on the Seurat R package^29^ integration algorithm to perform atlas-level data integration with millions of cells. The integration framework consisted of three major steps to align two or more data sets onto the same space: (1) using dimension reduction to derive embedding of the datasets in the same space; (2) using canonical correlation analysis (CCA) to capture the shared variance across cells between datasets and find anchors as mutual nearest neighbours between the two datasets; (3) aligning the low-dimensional representation of the datasets together with the anchors.

Integration and annotation of snmC-seq and snm3C-seq datasets.

For integration of the snmC-seq and snm3C-seq datasets, we utilized non-overlapping 100-kb chromosome bins as features. Our integration approach began with the selection of cluster-enriched features (CEFs)^124^ from both mC and m3C cells independently, based on their respective Leiden cluster assignments following methylome-based clustering described in previous sections. The CEF selection was implemented through the cluster_enriched_features function in the ALLCools package. For subsequent integration, we selected only CEFs that overlapped between the two datasets. We then constructed a principal component analysis (PCA) model using the mC cells and applied this model to transform the m3C cells. To prevent the embedding from being dominated by the initial principal components, we normalized each PC by its corresponding singular value.

To find anchors between mC and m3C cells, we first standardized the mC and m3C matrices of CEFs across cells using Z-score scaling. The resulting matrices were designated as X (mC:cell-by-CEF) and Yc(m3C:cell-by-CEF). We applied Canonical Correlation Analysis (CCA) to identify low-dimensional embeddings that captured shared variation between the two datasets. This was computed through single value decomposition of their dot products, expressed as 𝑈𝑆𝑉 = 𝑋𝑌 . We normalized the U and V matrices by dividing each row by its L2-norm. These normalized matrices were then used to identify five mutual nearest neighbors as anchors and to score these anchors using methodology consistent with the Seurat approach. To manage computational demands, we randomly selected 100,000 cells from each dataset as a reference to fit the CCA model. We then transformed the remaining cells onto the same canonical correlation space, using this fitted model. The canonical correlation vectors (CCVs) of 𝑋_𝑟𝑒𝑓_, and 𝑌_𝑟𝑒𝑓_ (denoted as 𝑈_𝑟𝑒𝑓_, 𝑉_𝑟𝑒𝑓_ respectively) were computed using 𝑈_𝑟𝑒𝑓_𝑆𝑉*^T^*_𝑟𝑒𝑓_ = *X*_𝑟𝑒𝑓_*Y^T^*_𝑟𝑒𝑓_ Subsequently, the CCVs for 𝑋_𝑞𝑟𝑦_ and 𝑌_𝑞𝑟𝑦_ (𝑈_𝑞𝑟𝑦_, 𝑉_𝑞𝑟𝑦_ respectively) were calculated using 𝑈_𝑞𝑟𝑦_ = 𝑋_𝑞𝑟𝑦_(𝑌*^T^*_𝑟𝑒𝑓_ 𝑉_𝑟𝑒𝑓_)/𝑆 and 𝑉_𝑞𝑟𝑦_ = 𝑌_𝑞𝑟𝑦_(X*^T^*_𝑟𝑒𝑓_ *U*_𝑟𝑒𝑓_)/𝑆. . For anchor identification, we concatenated the embeddings from both reference and query cells. Using the identified anchors, our integration projects the PCs of the query dataset (scm3C-seq), onto the PCs of the reference dataset (snmC-seq), while keeping the PCs of the reference dataset unchanged. This considerably reduced the time and memory consumption for computing the corrected high-dimensional matrix. Our integration approach utilized these anchors to project the principal components of the query dataset (scm3C-seq) onto those of the reference dataset (snmC-seq), while preserving the original principal components of the reference dataset. This strategic approach significantly reduced computational demands in terms of both processing time and memory requirements when generating the corrected high-dimensional matrix.

Following two rounds of iterative integration, we identified 59 co-clusters during the first iteration and ultimately derived a total of 342 co-clusters after completing the second integration round. The 2-month samples in the snmC-seq dataset originated from the BICCN whole mouse brain dataset^27^, which had previously undergone comprehensive annotation. For level 1 co-clusters, we applied a straightforward annotation rule: if more than 80% of the 2-month-old mC cells within a co-cluster shared the same subclass label, we assigned that label to the entire co-cluster. For co-clusters not meeting this criterion, we based annotations on the cluster information obtained after the second integration round, assigning labels according to the predominant BICCN cell subclass annotation within each cluster. To increase pseudo-bulk coverage for downstream analyses, we merged certain cell subclasses with similar functions and anatomical locations into unified major cell types. Further details of this process are provided in Table S6.

Integration of snmC-seq, snm3C-seq, snATAC-seq and 10X Multiome.

For our integrative analysis, we combined ATAC signals from snATAC-seq and 10X Multiome, and used methylation information from snmC-seq and snm3C-seq. We established snmC-seq data as our reference dataset, with the three other datasets serving as query datasets. PCA on gene-body signals alone was inadequate for capturing the full spectrum of open chromatin heterogeneity present in snATAC-seq data. Previous research^29^ has shown that Latent Semantic Indexing (LSI) applied to binarized cell-by-5kb bin matrices produces effective embeddings and clustering for snATAC-seq data. Following the previously introduced framework, we implemented this approach using binary sparse cell-by-5kb bin matrices to represent open chromatin regions.

We processed snmC-seq data by creating a cell-by-5kb bin matrix and calculating hypomethylation scores using a binomial distribution model, then binarized the matrix with a 0.9 threshold to produce a sparse binary matrix. Similar processing was applied to snATAC-seq data, where 1 represented at least one read detected in a 5 kb bin in a cell. Dimensionality reduction was performed using LSI with log term frequency and TF-IDF transformation to compute embeddings for both datasets. We then applied CCA with a downsampling framework of 100,000 cells from each dataset as the reference to align the methylation and chromatin accessibility data through anchor identification. Details in ATAC and methylation have been described in our previous work^27^. Our code can be found in the “Data and Code Availability” section.

### Cluster-level DNA methylome analysis

#### Age-Differentially DMG analysis

To identify Age-Differentially Methylated Genes (DMGs), we employed a reference genome comprising protein-coding and long non-coding RNA genes from the mouse GENCODE vm22 annotation, focusing on gene body regions. We compared methylation patterns across three age groups using single-cell level methylation fractions for both mCH and mCG contexts. For statistical analysis, we applied One-way ANOVA to identify genes with significant methylation changes, followed by multiple testing corrections using the Benjamini–Hochberg procedure to adjust p-values. We defined Age-related DMGs based on two criteria: 1) the gene was also identified as an age-differentially expressed gene (age-DEG) in our companion 10X Multiome dataset (FDR < 0.05, log2[18mo/2mo] > 0.1); and 2) the gene showed significant methylation differences in our ANOVA results (FDR < 0.05). The complete analytical pipeline and implementation details are available in our “Data and Code Availability” section.

#### Age-Differentially DMR analysis

Following clustering analysis, we merged individual cell ALLC files into pseudo-bulk profiles using the allcools.merge-allc command, grouping cells by cell-type and age. Because male and female samples were processed using different sequencing methods, we focused on identifying methylation changes common to both sexes to minimize technical variation. For each cell-type-age combination, we merged pseudo-bulk ALLC files from both sexes, normalizing them to ensure equivalent total cytosine coverage. Using these sex-normalized pseudo-bulk ALLC files, we applied a multi-factor model through DMLfit.multiFactor from DSS (version 2.42.0)^125^ package, with a design matrix incorporating both cell type, replicate, and age as variables. We then tested specifically for age-related differential methylation sites (DMS) using DMLtest.multiFactor, which implements a beta-binomial regression model optimized for detecting differential methylation signals in methylation datasets. To identify differentially methylated regions (DMRs), we employed the callDMR function and applied rigorous selection criteria, including: (1) p-value < 0.01; (2) A minimum region span of 1 base pair containing at least 1 CpG site; (3) A requirement that at least 50% of CpGs within each candidate region demonstrate significant differential methylation. Additionally, we merged adjacent DMRs when they were separated by less than 50 base pairs. The complete analytical workflow and implementation details are available in our “Data and Code Availability” section.

#### Transcription Factor motif enrichment analysis

To identify enriched transcription factor (TF) binding motifs, we employed HOMER^39^ software, applying the FindMotifsGenome.pl function to analyze the ±200bp regions corresponding to each age-associated differentially methylated region (age-DMR). We used whole-genome sequences as the background reference set. Motifs from the HOMER known motifs database were considered significantly enriched when the false discovery rate was below 5% (q < 0.05), unless specified otherwise.

For visualization purposes, we applied different filtering criteria to the enrichment results. In heatmaps displaying age-hypomethylated DMR enrichment patterns (Figure 2D), we included only motifs that met all of the following conditions: (1) enrichment p-value < 10^-30^, (2) log2[% of Target Sequences with Motif/% of Background Sequences with Motif] > 1.1, and (3) significant enrichment in at least three cell types. For age-hypermethylated DMR enrichment visualization, we applied less stringent criteria, including motifs with: (1) p-value < 10^-10^, (2) fold enrichment (log2 [% of Target Sequences with Motif/% of Background Sequences with Motif]) greater than 1.1, and (3) significant enrichment in at least one cell type.

#### Methylation Hotspot analysis

We segmented the mm10 genome into non-overlapping 500 base pair (bp) chunks using “bedtools makewindows” command. Then we selected significant age-related differentially methylated sites (age-DMSs) using a statistical threshold of p < 0.01 using results from DMLtest.multiFactor introduced in the previous section. These significant sites were then intersected with both hypomethylated (age-hypo) and hypermethylated (age-hyper) DMRs, retaining only the DMSs that overlapped with these regions. We subsequently aggregated all cell-type-specific age-associated DMSs across the different cell types examined using “bedtools intersect” command. For each 500 bp genomic chunk, we quantified the total number of age-associated DMSs present across all cell types. Chunks containing more than 10 age-associated DMSs were classified as methylation “hotspots”.

#### GAT analysis

To assess the association between methylation hotspots and transposable element (TE) subfamilies, we employed the Genomic Association Tester (GAT) package (v1.3.6), specifically using the gat-run.py function. This statistical framework evaluated whether the identified hotspot regions were significantly enriched for specific TE subfamilies compared to the genomic background. The analysis involved 10,001 permutations, where the algorithm randomly shuffled segments across the whole genome to generate a null distribution of expected overlaps. This permutation-based approach allowed us to determine if the observed overlaps between hotspots and TE subfamily regions occurred at frequencies significantly higher than expected by chance. We established stringent criteria to define significant enrichment: a false discovery rate below 1% (q-value < 0.01) and a fold change (target/background) > 1.1. These thresholds ensured that only robust and biologically meaningful associations between methylation hotspots and TE subfamilies were reported.

### Transposon Element Clustering

Transposable element (TE) annotations were obtained from RepeatMasker (https://www.repeatmasker.org/). To analyze methylation patterns across TEs, we constructed a matrix representing methylation levels for each TE region across all cell types. Due to the large number of TE regions and computational constraints, we simplified the analysis by binarizing the methylation matrix: regions with methylated mCG fractions < 0.75 were assigned a value of 1 (hypomethylated), while regions with mCG fractions ≥ 0.75 were assigned a value of 0 (not hypomethylated). From this binarized matrix, we selected TE regions that exhibited hypomethylation in at least one cell type for further analysis for downstream clustering. Dimensional reduction was performed on this sparse binary matrix using the ALLCools.clustering.lsi function. Subsequent clustering analysis was conducted using the Scanpy package. The complete computational workflow, including detailed code implementation, is available in the “Data and Code Availability” section.

### Chromatin conformation analysis

The chromatin conformation was examined at both individual cell and aggregated levels. For individual cell analysis, we processed data following completion of snm3C-seq mapping and quality filtering procedures outlined in earlier sections. Subsequently, we excluded genomic segments that coincided with the ENCODE blacklist (version 2)^118^ using “hicluster filter-contact” command. We constructed individual cell chromatin interaction matrices using long-range cis contacts (contact anchor distances > 2,500 bp) and trans contacts at three distinct resolutions: 100-kb for analyzing chromatin compartmentalization; 25-kb for evaluating domain boundary features; and 10-kb for chromatin loop and enhancer analysis. Contact matrix imputation was performed using the scHiCluster^113^ package (version 1.3.5) using “hicluster prepare-impute” command. This imputation methodology first applied Gaussian convolution (pad = 1) and subsequently employed a random walk with restart algorithm on the convoluted matrix. For 100-kb matrices, imputation was conducted across whole chromosomes, while for 25-kb matrices, imputation was confined to contacts within 10.05 Mb. At 10-kb resolution, imputation was further restricted to contacts within 5.05 Mb. For pseudo-bulk analysis, we merged cells by group (categorized by cell-type, cell-type-age, or cell-type-age-replica). We used the “hicluster merge-cell-raw” command to sum raw matrices or the “hicluster merge-cool” command to calculate the mean of imputed matrices. To maintain analytical rigor in all 3C-related age-differential analyses, we established consistent sampling parameters across groups. We ensured the total number of nuclei between groups is the same and required at least 100 nuclei per group to guarantee sufficient genomic coverage.

#### Contact distance distribution analysis

Contact distance distributions for individual cells were generated utilizing the “hicluster contact-distance” command. This created histograms based on the spatial separation between contact anchor pairs. We employed logarithmic binning on a log2 distance scale using 0.125 as the step increment, with the full range extending from 2500 bp to 249 Mb (corresponding to the longest chromosome in the genome). Each histogram bin (denoted as the i-th bin) contained the count of contacts whose distances fell between 2500 × 2^(i×0.125)^ and 2500 × 2^((i+1)×0.125)^. As presented in Figure S10A, we calculated the short-long ratio by dividing the proportion of contacts in bins 51-76 (representing distances from 200kb to 2 Mb) by the proportion found in bins 103-114 (representing distances from 20 Mb to 50 Mb).

#### Compartments Analysis

For compartment analysis, we employed cell-type-specific pseudo-bulk raw contact matrices for each chromosome at 100 kb resolution. We implemented filtering to exclude 100-kb bins exhibiting atypical coverage patterns. Specifically, we defined the coverage of bin i on chromosome c (represented as Rc,i) as the sum of values in the i-th row of the chromosome c contact matrix. We retained only those bins with coverage values falling between the 99th percentile of Rc and a threshold calculated as twice the median of Rc minus the 99th percentile of Rc. Following selection, we normalized the contact matrices by distance and calculated Pearson’s correlation matrices from these normalized datasets^62^. These consolidated contact maps served as input for principal component analysis (PCA) models constructed for individual chromosomes. We utilized the first principal components (PC1) as compartment score indicators, adjusting the model sign to ensure compartments with higher CpG density displayed positive scores. To validate our approach, we visually examined the PC1 values from the consolidated matrices, confirming their correspondence to the characteristic plaid pattern observed in correlation matrices rather than chromosome arm effects.

To ensure each age group is transformed with the same PCA model, the raw contact matrices of each age group within the same cell types are filtered and converted to the correlation matrices in the same way as described above, and were then transformed with the PCA models. We used the merged raw matrices for fitting the PCA model, and transformed the correlation matrices of raw matrices in each cell type as raw compartment scores. Differential compartments (DCs) were identified with dcHiC ^126^ between all age groups within each cell type using the raw compartment scores as input. Age-related differential compartments were identified as differential with a traditional q-value < 0.05.

We calculated saddle plots and quantified compartment strengths following the previous method^127^. For each individual chromosome, we arranged all 100kb bins in rank order based on their compartment scores and subdivided them into 50 groups of equal intervals. The distance-normalized interaction strength between each pair of bins were averaged within each group. In the visualization framework, the axes are ranked by the compartment score of the groups so that BB (B compartment to B compartment) interactions are on the top left and AA (A compartment to A compartment) interactions are on the bottom right.

#### Domain boundary analysis

We identified domains and calculated insulation scores with scHiCluster at 25 kb resolution using “hicluster domain” command. Domain identification for individual cells was performed by applying the TopDom algorithm^128^ to the imputed matrices at 25 kb resolution. For each bin, we calculated insulation scores across each cell-type-age group using pseudo-bulk imputed matrices (derived by averaging across individual cells) and a window size of 10 bins. We defined boundary probability for a specific bin as the fraction of cells within a group where that bin was identified as a domain boundary. To detect differential domain boundaries among the three age groups within each cell type, we constructed a 3×2 contingency table for each 25kb bin, with rows representing the count of cells from each group where the bin was either classified as a boundary or not. Statistical analysis involved calculating the Chi-square statistic and corresponding P value for each bin, with significant differential boundary bins identified using a threshold of FDR < 1 × 10^-3^. Additionally, we imposed a requirement that the difference between the maximum and minimum boundary probability is larger than 0.05. In Figure 4D-4F, the CTCF ChIP-seq data from mouse forebrain tissue is obtained from the ENCODE database (https://www.encodeproject.org/experiments/ENCSR677HXC/).

#### Boundary-proximal genes

For each age-DB, we applied boundary information generated using the “hicluster domain” command in scHiCluster ^113^, following the same methodology used for the single-cell files described in the above sections. Briefly, this approach processes pseudobulk-level imputed contact files at 25 kb resolution which are grouped by their cell type and age. For each age-DB, we identified the nearest neighboring boundary in the 2-month-old group. We then intersected genes located within these regions using the GENCODE vM22 GTF file, identifying a set we term “boundary-proximal genes. Detailed code is in the “Data and Code Availability” section.

#### Identification of loops and differential loops

Chromatin loop detection was performed as previously described^129^. Briefly, we identified chromatin loops using the schicluster.loop.call_loop function for each cell-type-age or cell-type-region-age group. Chromatin loop identification was conducted within the 50 kb to 5 Mb range. The process involved transforming imputed chromosome matrices (Q) through log-transformation and Z-score normalization at each diagonal (yielding E), followed by subtraction of a local background between ≥30 kb and ≤50 kb (producing T). Statistical significance was established using t-statistics to quantify how significantly E and T deviated from zero across cells. Differential loops were identified in two comparisons: between all age groups within the same cell type, and between age groups within cell-type-region categories. To compare loop interaction strength between different cell groups, we performed ANOVA to compute the statistical difference for each loop identified in at least one age group. Age-differential loops were defined as the top 15% of loops based on ANOVA values. Details can be found in the “Data and Code Availability” section.

#### Activity by Contact analysis

We employed the Activity-by-Contact (ABC) model^130^ to identify putative enhancers and their target genes in each group. Briefly, the ABC model integrates pseudo-bulk raw contact frequencies from 3C data with enhancer activity measurements to predict enhancer-gene pairs.

We identified candidate cis-regulatory elements (cCREs) using two methods: (1) ATAC-seq peaks from the corresponding cell-type-age group, and (2) cell-type-specific hypomethylated regions (hypo-DMRs; mCG fraction < 0.7) from the corresponding cell type^27^. Enhancer activity was quantified using RPKM values for ATAC-seq data and (1 - mCG fraction) for methylation data. Predictions with an ABC score ≥ 0.02 were classified as positive interactions and used for downstream analysis. Detailed code can be found in the “Data and Code Availability” section.

## MERFISH experiments

### MERFISH gene panel design

Gene selection criteria were applied to design a panel of 500 genes, consisting of 233 cell type marker genes and 267 genes related to aging biology. Genes in the GENCODE vm22 GTF file were filtered based on length parameters (>1 kb and <1000 kb). For cell type marker genes, we utilized the GenePanelDesign package (https://github.com/jksr/GenePanelDesign). We employed Welch’s t-test to identify differentially expressed marker genes among major cell types, selecting those with FDR < 0.05 and ranking them by standardized mean difference, calculated as 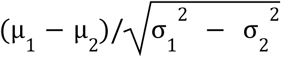, where µ and σ represent the mean value and standard error of each group, respectively. We also manually added known cell type markers from previous studies. All markers were validated to ensure robust expression in the RNA dataset with sufficient diversity to distinguish cell clusters. We confirmed that these markers effectively separated both RNA and mC cells by cell type. The remaining 267 genes were selected based on specific criteria: (1) methylation-related genes (e.g., Dnmt3, Tet1); (2) age-associated differentially methylated genes in Figure S5; (3) Alzheimer’s disease-related genes (e.g., Trem2, Nlrp3, Il18); (4) senescence-associated secretory phenotype (SASP) genes (e.g., Atf3, Irf1); (5) hotspot genes from our previous analyses (e.g., Pax5, Sgk1); (6) immune cell markers (e.g., Ptprc, Cd3d, Cd69, Cd19); (7) transcription factors with regulatory roles for hotspot genes; and (8) positive control genes from previous brain aging studies ^32^ (e.g., Onecut2, Pakap). Encoding probes for the final 500-gene panel were designed and synthesized by Vizgen (Table S4).

### MERFISH tissue preparation and imaging

Mouse brains were harvested intact and sliced coronally at the chosen sections. Individual slices were placed in OCT compound, rapidly frozen in isopentane and dry ice, and maintained at −80°C for subsequent processing. Using a Leica CM1950 cryostat, the OCT-embedded tissues were cut into 12-μm-thick coronal sections. These sections underwent immediate fixation in pre-warmed (37°C) 4% formalin for 30 minutes, followed by permeabilization with 70% ethanol according to the manufacturer’s guidelines. The Vizgen sample preparation kit (10400012) was utilized for all subsequent processing, including probe hybridization and gel embedding, following the standard protocol provided by the manufacturer. Imaging of each tissue section was performed on a MERSCOPE instrument using the MERSCOPE 500 Gene Imaging Kit (Vizgen, 10400006).

### MERFISH data preprocessing

MERFISH data processing and quality control were performed using MERSCOPE software (version 2023-01), which handled both image analysis and transcript detection. For cell segmentation, we employed the ’nuclei’ model integrated within the cellpose (v2.1.0) package. The resulting cell-by-gene expression matrix was constructed by aggregating all detected transcripts within the boundaries of each identified cell. We implemented a multi-parameter quality filtering approach to eliminate aberrant cellular objects from the expression matrices, similar to [https://www.nature.com/articles/s41586-024-08334-8]. Specifically, we excluded cells based on the following parameters: (1) cellular volume > 3× the median volume (μm^3^) or less than 100 μm^3^; (2) total transcript counts < 20 or > 4000; (3) volume-normalized transcript abundance < the 2nd or > the 98th population percentile; (4) < 5 distinct genes detected; and (5) presence of > 5 negative control (blank genes) probes.

### Integration between MERFISH and snRNA for annotation

Following the quality control step, the remaining MERFISH dataset was integrated with the snRNA dataset using the previously described ALLCools integration framework. In brief, we utilized the well-annotated BICCN scRNA-seq dataset^31^. For each MERFISH sample, we selected the snRNA cells located in the brain regions included within that sample. To perform cluster-level annotation, we first conducted two rounds of iterative clustering on the MERFISH samples. We then performed two rounds of integration between the MERFISH and snRNA dataset. During the first round of integration, if a co-cluster had 70% of RNA cells with the same annotated subclass label, we assigned that annotation to all the MERFISH cells in that co-cluster. Otherwise, we proceeded to the second round of integration, where we assigned the annotation if >80% of RNA cells in the cluster shared that annotation. For clusters that did not pass the 80% same annotation rule, we left them unannotated, as they might not have been included in the aging dataset and the gene markers in the panel were insufficient to distinguish them.

### MERFISH and age-DEG analysis

To identify differentially expressed genes (DEGs) with the MERFISH dataset, we grouped cells by their coronal slices and compared across age groups within each cell type. The identification of DEGs involved the following steps. We filtered cells with less than 10 genes expressed and genes with less than 5 cells expressed. For each sample, the data were normalized to total counts using scanpy.normalize_total() function and log-scaled using scanpy.pp.log1p(). We performed two types of statistical tests. First, we performed the Kruskal-Wallis test, a non-parametric alternative to ANOVA, to identify genes with significant expression differences across three different groups simultaneously. We also compared DEGs pairwise among three age groups, where we conducted pairwise comparisons using the Wilcoxon rank-sum test (implemented in the diff_ranksum function) for all possible sample group combinations. For both methods, the p-values were adjusted for multiple testing using the Benjamini-Hochberg false discovery rate (FDR) method, controlling for type I errors while maintaining statistical power.

### Machine learning analysis

#### Input Data generation

We generate a cell-type-specific multimodal dataset for machine learning applications. Here we introduce how we generated the input dataset, with detailed code in the “Data and Code Availability” section. Our model integrates three epigenetic modalities across seven features (𝑋) to predict age-related gene expression changes (𝑌). The input data generation process is described below:

𝑋:

For methylation modality, three different methylation features were extracted: 1) Genebody DMRs: For each gene, we collected all age-DMRs located within the gene body ± 2kb. Input features included DMR mCG methylation levels at 2mo, 9mo, and 18mo timepoints, gene length, distance to the transcription start site (TSS), DMR length, and additional relevant parameters. 2) Gene body mCG fraction: We calculated the average mCG fraction across the gene body ± 2kb at 2 month, 9 month, and 18 month timepoints. Input features included these methylation levels, gene length, and p-values from age-related differentially methylated gene (DMG) analysis. 3) Gene body mCH fraction: Identical parameters as mCG fraction analysis but using non-CpG (mCH) methylation levels instead.

Chromatin accessibility (ATAC) modality: ATAC peak information was compiled similarly to genebody DMRs. Values represent ATAC RPKM. We also included an annotation of whether each peak represented an age-related differentially accessible region (age-DAR).

For chromatin conformation (3C) modality, three chromatin conformation features were analyzed: 1) 3C chromatin loops: We utilized loop calling results as described in previous sections. For each gene body ± 2kb, we included all loops with either anchor overlapping this region. Input features included normalized contact frequencies at 2mo, 9mo, and 18mo timepoints, gene length, distance between loop anchors, distance between loop and TSS, and classification of age-related differential loops. 2) Enhancer ATAC peaks: We incorporated results from activity-by-contact analysis described in previous sections. 3) Enhancer hypo-DMRs: same as enhancer ATAC peaks, it has been described in the previous section.

𝑌:

Gene expression: DEG analysis was performed using MAST^131^. Genes were categorized into three distinct groups based on results from age-related differential expression analysis: 1) Age-upregulated DEGs: FDR < 0.01 and log2[18mo/2mo] > 0.3; 2) Non-DEGs: FDR > 0.01 or |log₂[18mo/2mo]| < 0.3; 3) Age-downregulated DEGs: FDR < 0.01 and log₂[18mo/2mo] < −0.3

For all transformer models, we balanced the dataset by including all DEGs (both up- and down-regulated) and then randomly sampling from the non-DEGs to achieve a count equal to 50% of the total DEG count. This approach ensured the non-DEG class did not overwhelm the DEG classes during model training. Data normalization was then performed across all features for each cell type using either z-score or min-max scaling.

### EAT model

We developed a transformer-based model, EpiAgingTransformer (EAT). The model starts with seven transformer modules, followed by a self-attention aggregator and a multiple-layer perceptron (MLP) classifier.

We used the sequence dataset (7 epigenetic feature matrices, where in each matrix the rows represent different genomic loci and the columns are the features associated with each loci) to train the EAT model. To ensure robust model evaluation, we implemented 5-fold cross-validation, training on four folds and predicting on the held-out fold, thereby generating predictions for the entire dataset without data leakage.

To handle the variable-length input data, in the EAT model, we employed seven transformer modules to extract meaningful information from different modalities and a self-attention aggregator to aggregate information across modalities into a unified gene embedding. For each gene, we processed the sequence of loci features within each modality through a transformer model, utilizing a Classify (CLS) token embedding to condense loci-specific information into a single modality embedding vector. Subsequently, we aggregated information from all seven modality embeddings 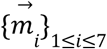 using the self attention^86^. The self-attention aggregator computed attention weights, 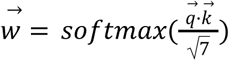, to determine each modality’s contribution to the final gene representation, where 𝑞 and 𝑘 are transformed vectors computed from the modality 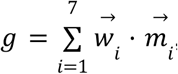. The final gene representation was computed as, [’inline], applying a weighted summation of modality embeddings based on their attention weights. The resulting gene embedding was then processed through an MLP classifier to predict age-related differential expression patterns, categorizing genes as upregulated, downregulated, or unchanged.

### Logistic regression model for classification

To transform the dataset to have a predefined number of features across all genes, we aggregate across all features within each group. We used mean, median, and standard deviation for each feature where applicable. We term this dataset the ‘block dataset’ below. A Logistic Classifier on the block dataset was implemented using the sklearn.linear_model.LogisticRegression class from sklearn (v1.5.1). For each cell type, we performed hyperparameter tuning using sklearn GridSearchCV on 5 splits, and then trained the model on the same splits using the generated hyperparameters to generate predictions for all genes in our dataset. Confusion matrices were generated using sklearn.metrics.confusion_matrix and normalized using norm=”pred”. The True Positive (TP) rate per label was then extracted for downstream analysis. Detailed code in the “Data and Code Availability” section.

### XGBoost model

A Gradient Boosted Tree classifier was trained using XGBoost’s Python package (v2.1.2). First, all features with more than 50% missing values in a given cell type were dropped from the block dataset. The model was then trained in successive rounds of feature selection, whereby in each round of training, hyperparameter tuning was performed using hyperopt (v0.2.7), and feature importance rankings for each classification task were generated using SHAP (v0.46.0). Briefly, SHAP is a Python package made to calculate Shapley^132^ values, which are game-theory-based axiomatic measurements of a feature’s contribution toward a specific model prediction. After features were ranked using their Shapley values, only the top 50% of features from each classification task were kept for the next iteration. Confusion matrices were generated in the same way as for the Logistic Classifier model. The difference between the TP rate of the XGBoost model and the Logistic Classifier was calculated and a one-sided t-test for difference from 0 was performed using scipy (v1.13.1). The detailed code is in the “Data and Code Availability” section.

## Supplementary tables

Table S1. Brain Dissection Region Annotation

Table S2. Metadata for snmC-seq3

Table S3. Metadata for snm3C-seq

Table S4. MERFISH gene panel

Table S5. Metadata for MERFISH

Table S6. Major Cell Type Annotation Table

---

**Figure S1.**
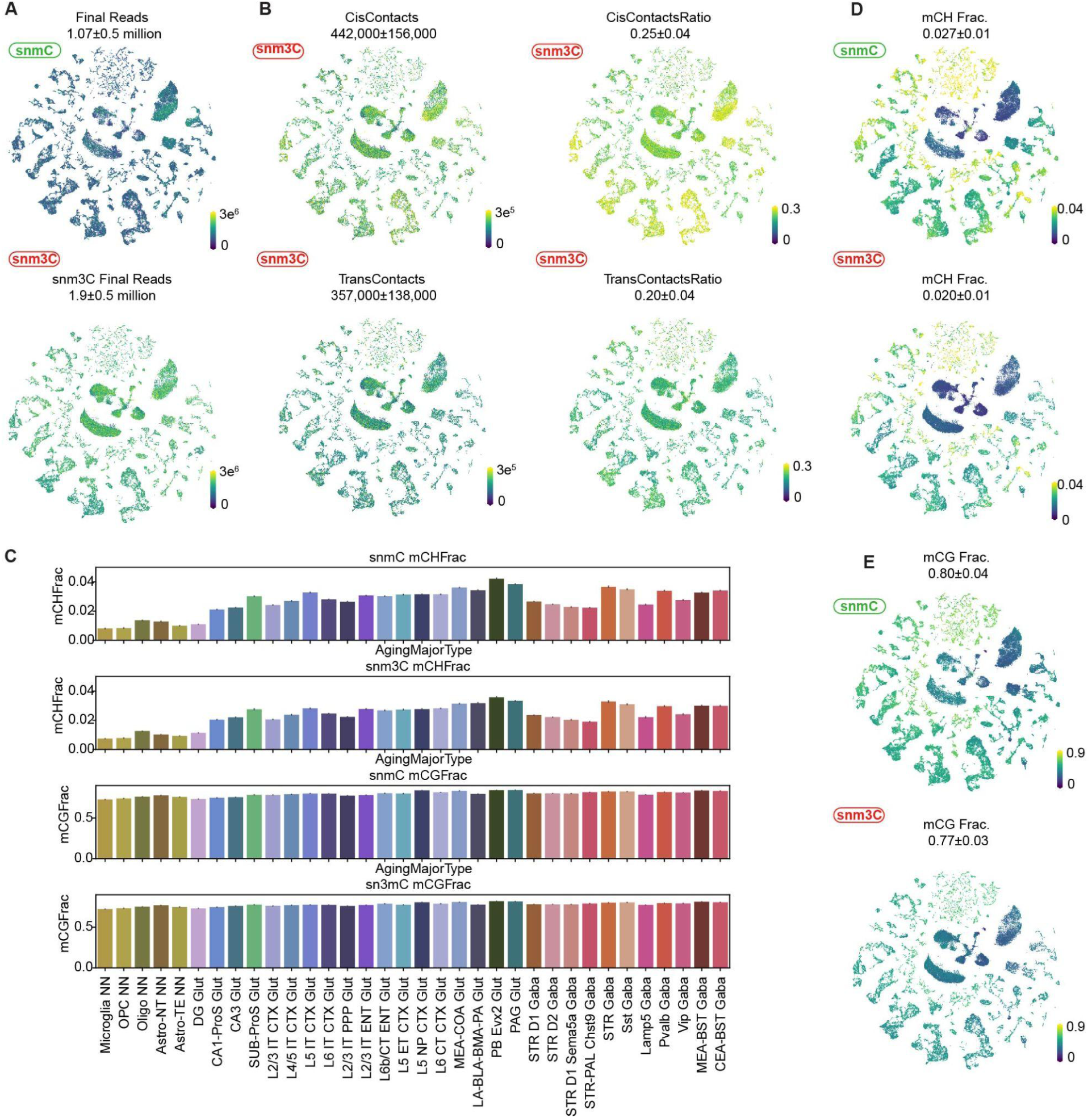
Quality control analysis for snmC and snm3C datasets. **(A)** The number of final pass QC reads in snmC-seq (top) and snm3C-seq (bottom), depicted by t-SNE. **(B)** The number (left) and ratio (right) of cis-long and trans contacts in snm3C-seq, depicted by t-SNE. **(C)** Barplots exhibit the average global mCH (top 2 rows) and mCG (bottom 2 rows) fractions for all major cell types. Bars are colored by cell types and cell types are ordered by the global mCG level. **(D-E)** The global mCH **(D)** fraction and mCG **(E)** fraction for snmC-seq (top) and snm3C-seq (bottom), depicted by t-SNE.

**Figure S2.**
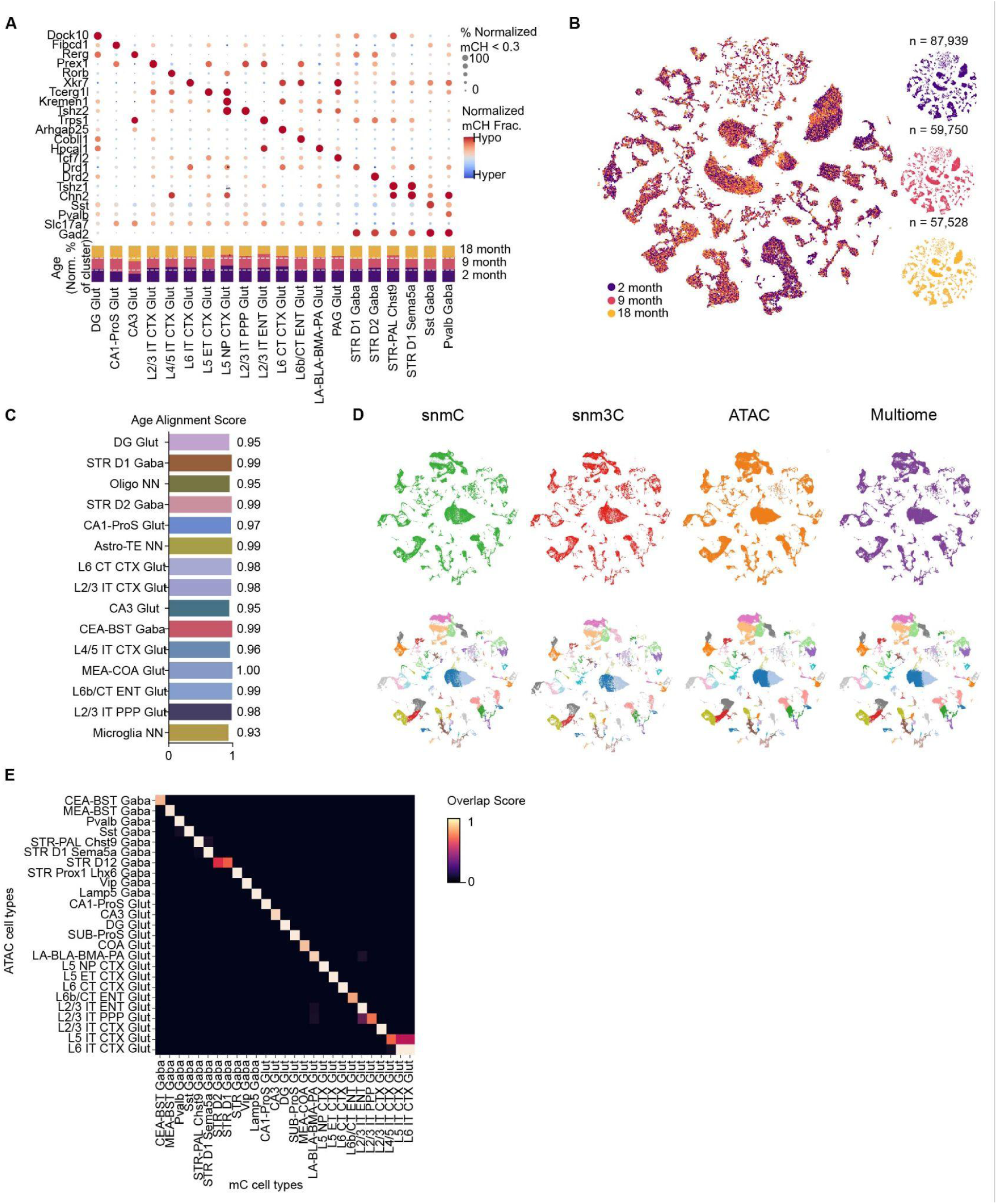
Cross-modality integration of snmC-seq, snm3C-seq, snATAC-seq, and 10X Multiome data. **(A)** Top: dot plot shows marker genes’ average mCH methylation level across major cell types; Color represents the genebody mCH level, size demonstrates the ratio of cells with mCH level < 0.3. Bottom: barplot represents the ratio of age group distribution in each cell type. **(B)** t-SNE visualization of integrated snmC-seq and snm3C-seq datasets, with cells colored by age group. **(C)** Barplot displays the alignment scores of age groups among each major cell type, calculated from the snmC-seq and snm3C-seq integration. Scores range from 0 (no alignment) to 1 (complete alignment). **(D)** t-SNE visualization of neuronal populations across four modalities (snmC-seq, snm3C-seq, snATAC-seq, and 10X Multiome). Cells are colored by modality (top) and Leiden clustering (bottom), with cells from other modalities shown in light grey. **(E)** Confusion matrix heatmap shows cell type correspondence between snmC-seq and snATAC-seq datasets after cross-modality integration, quantified by cluster overlap scores^28,102^.

**Figure S3.**
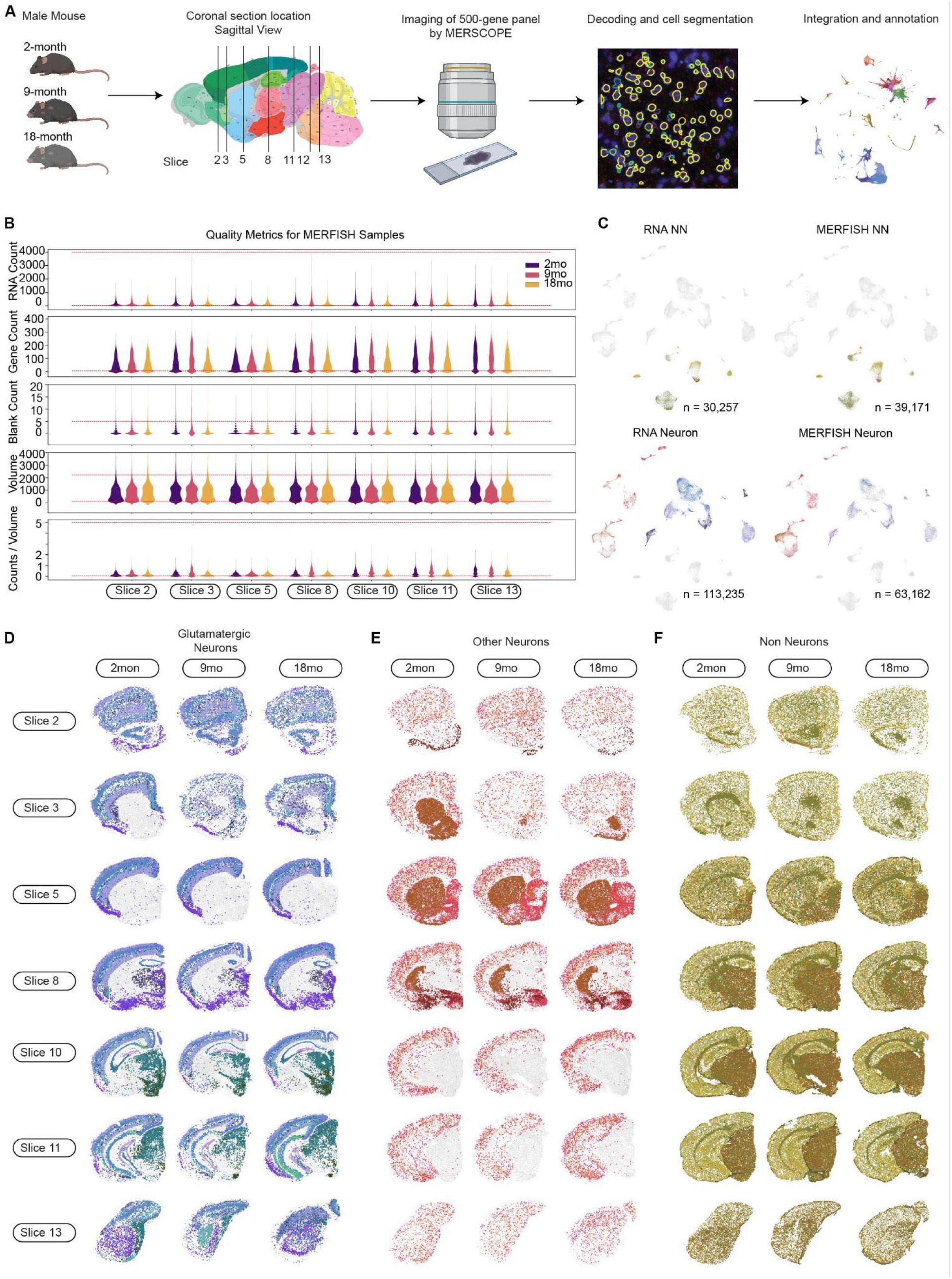
MERFISH data processing and cell type annotation. **(A)** Schematic workflow of MERFISH data generation and analysis, including sample age selection, tissue slice preparation, imaging procedures, and computational analysis pipeline. **(B)** Quality control metrics across MERFISH samples showing RNA total counts, gene counts, blank gene detection, cell volume (μm^3^), and RNA density (counts/volume). Red lines indicate quality filtering thresholds. **(C)** t-SNE plot of integrated MERFISH (right) and scRNA (left) dataset colored by cell types. **(D-F)** Spatial distribution of MERFISH cells across age groups (2-, 9-, and 18-months, left to right), annotated through scRNA-seq integration (Methods). Cell populations shown: glutamatergic neurons **(D)**, other neuronal cell types **(E)**, and non-neuronal cells **(F)**.

**Figure S4.**
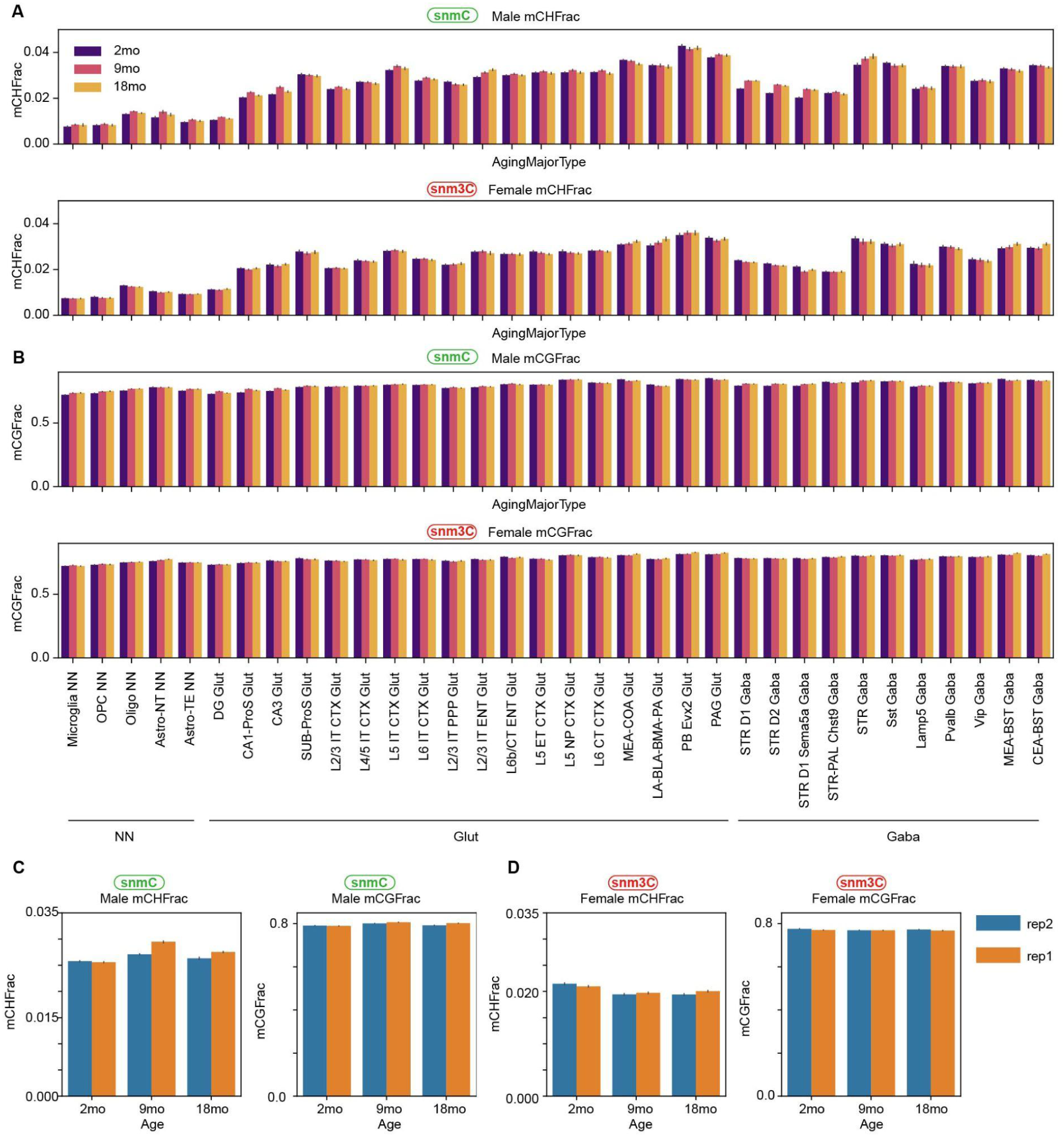
Modest global DNA methylation changes during aging. **(A-B)** Global mCH **(A)** and mCG **(B)** methylation levels across age groups among major cell types, measured by snmC-seq (top) and snm3C-seq (bottom). Bars are colored by age groups. **(C-D)** Barplots show global mCH (left) and mCG (right) levels of age groups among replicates in snmC-seq **(C)** and snm3C-seq **(D)**.

**Figure S5.**
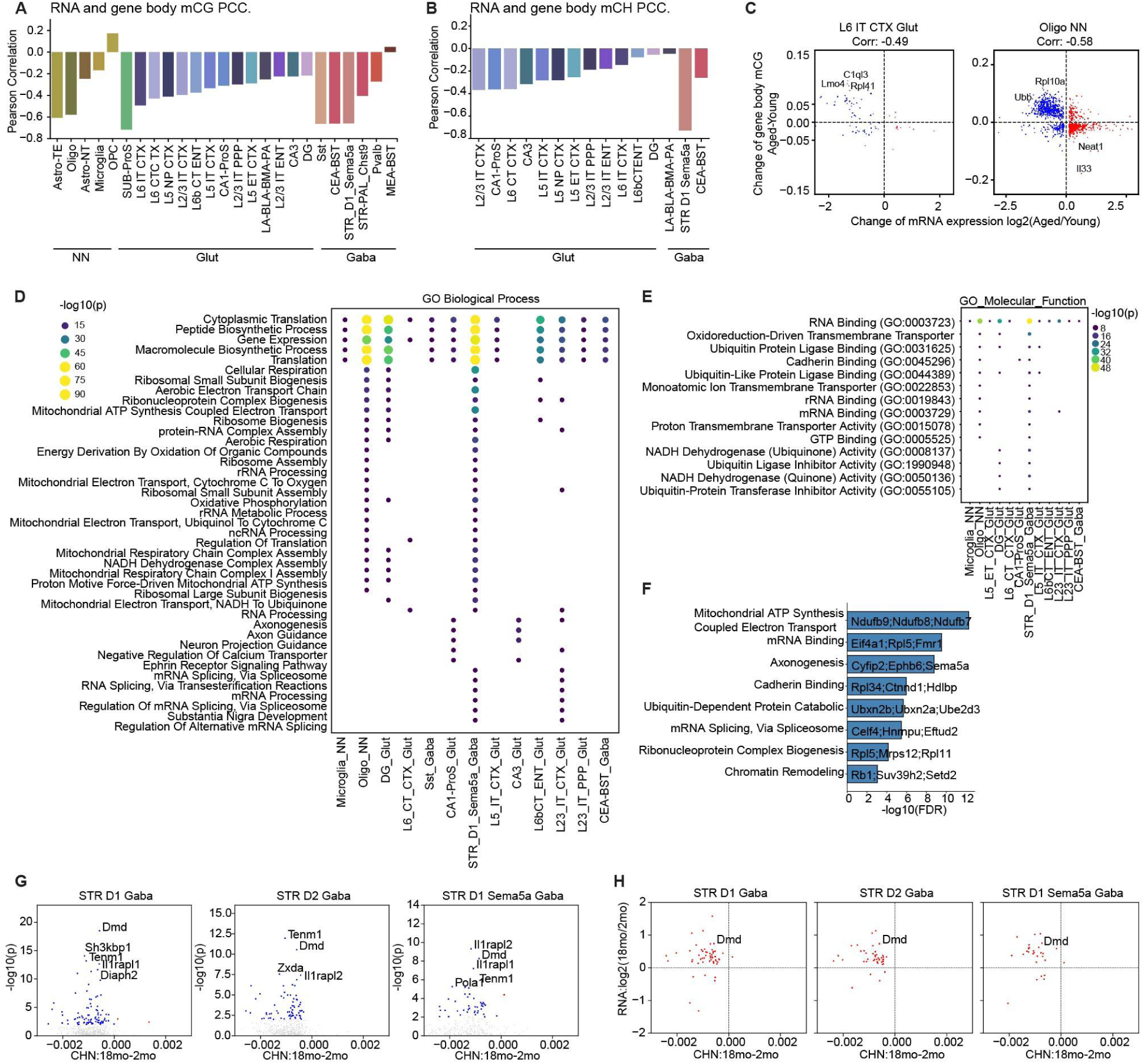
Age-related DMGs and associated biological processes. **(A-B)** Pearson correlation coefficients between age-related methylation changes (18mo - 2mo) and gene expression changes (log2[18mo/2mo]) across cell types, showing **(A)** CG (ΔmCG) and **(B)** non-CG (ΔmCH) methylation in gene bodies. Bars are colored by cell types. **(C)** Scatterplot shows the correlation between gene expression changes (log2[18mo/2mo], x-axis) and gene body CG methylation changes (ΔmCG: 18mo - 2mo, y-axis) in “L6 IT CTX Glut” and “Oligo NN”. Each dot represents an age-related DMG. **(D-E)** Enrichment of **(D)** GO biological processes, **(E)** GO molecular functions among age-related DMGs across cell types. Dot color and size indicate statistical significance (−log10[P]). **(F)** Barplot depicts the enriched GO processes for all age-related DMGs from all cell types, with specific DMGs annotated for each enriched pathway. **(G)** Scatterplot exhibits the mCH change (18mo - 2mo) on X chromosome genes across cell types **(H)** Scatterplot shows the correlation between mCH (18mo - 2mo) and RNA (log2[18mo/2mo]) change of X chromosome genes across cell types

**Figure S6.**
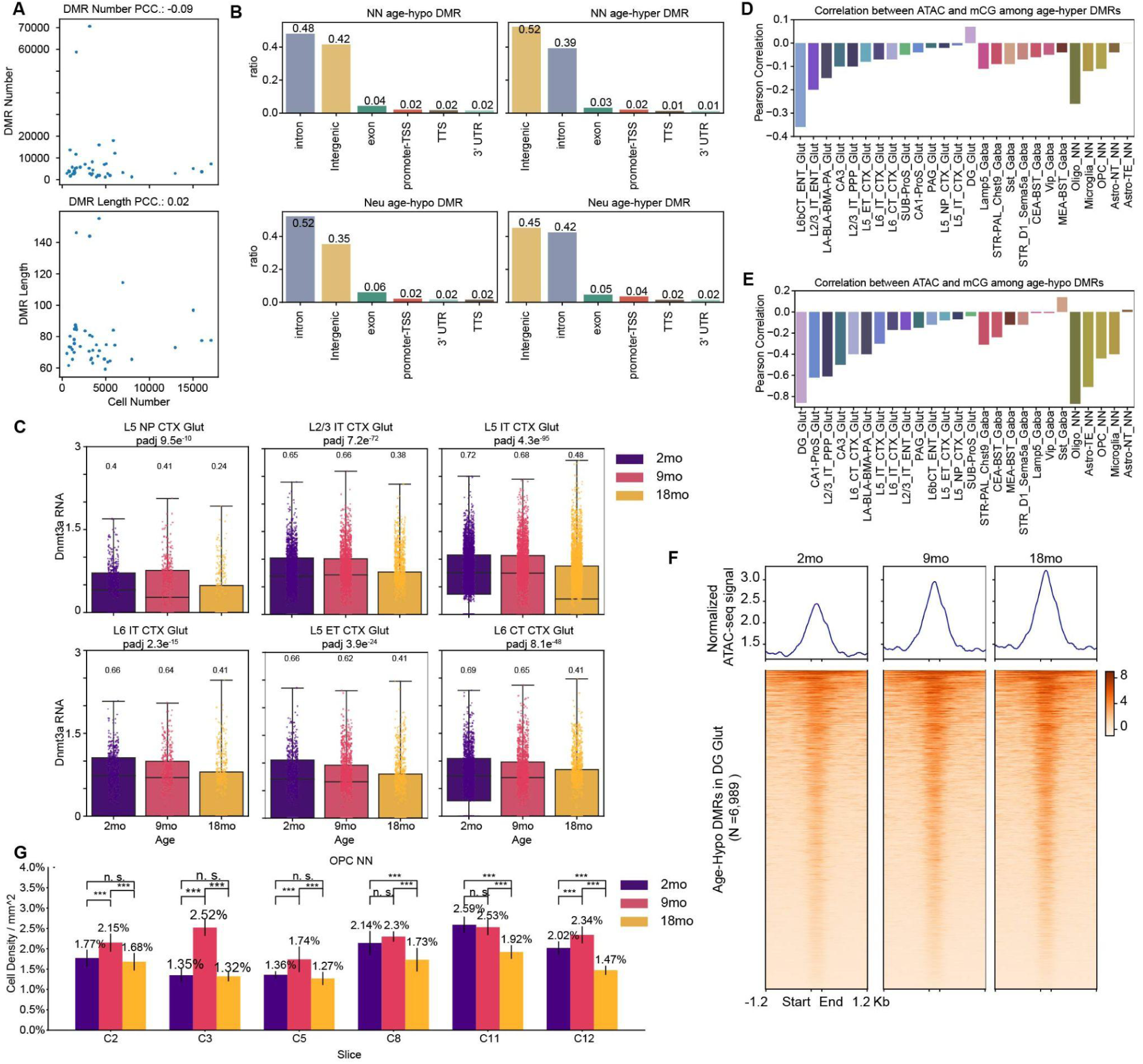
Analysis related to cell-type-specific age-related DMRs. **(A)** Scatter plot exhibits the relationship between total cell number and (top) number of DMRs, and (bottom) average DMR length. Each point represents a cell type. **(B)** Barplots depict the genomic distribution of age-related DMRs in non-neuronal (top) and neuronal (bottom) cells, separated by age-hypo (left) and age-hyper (right) DMRs. Numbers above each bar represent the ratio of age-DMRs in the corresponding genomic feature. **(C)** Boxplot shows RNA expression level of *Dnmt3a* across age groups among cell types. Each dot represents a cell, with color indicating age group. **(D-E)** Barplots show the Pearson correlation between ATAC RPKM (Log2[18mo/2mo]) and mCG (ΔmCG: 18mo - 2mo) changes of age-hyper **(D)** and age-hypo DMRs **(E)**. **(F)** Chromatin accessibility among age-hypo DMRs in “DG Glut”. Top: Distribution of ATAC-seq signal across DMRs. Bottom: Aggregated heatmap of normalized ATAC-seq signals across individual DMRs (rows). **(G)** Barplot shows the cell density of ‘OPC NN’ across MERFISH slices among different age groups.

**Figure S7.**
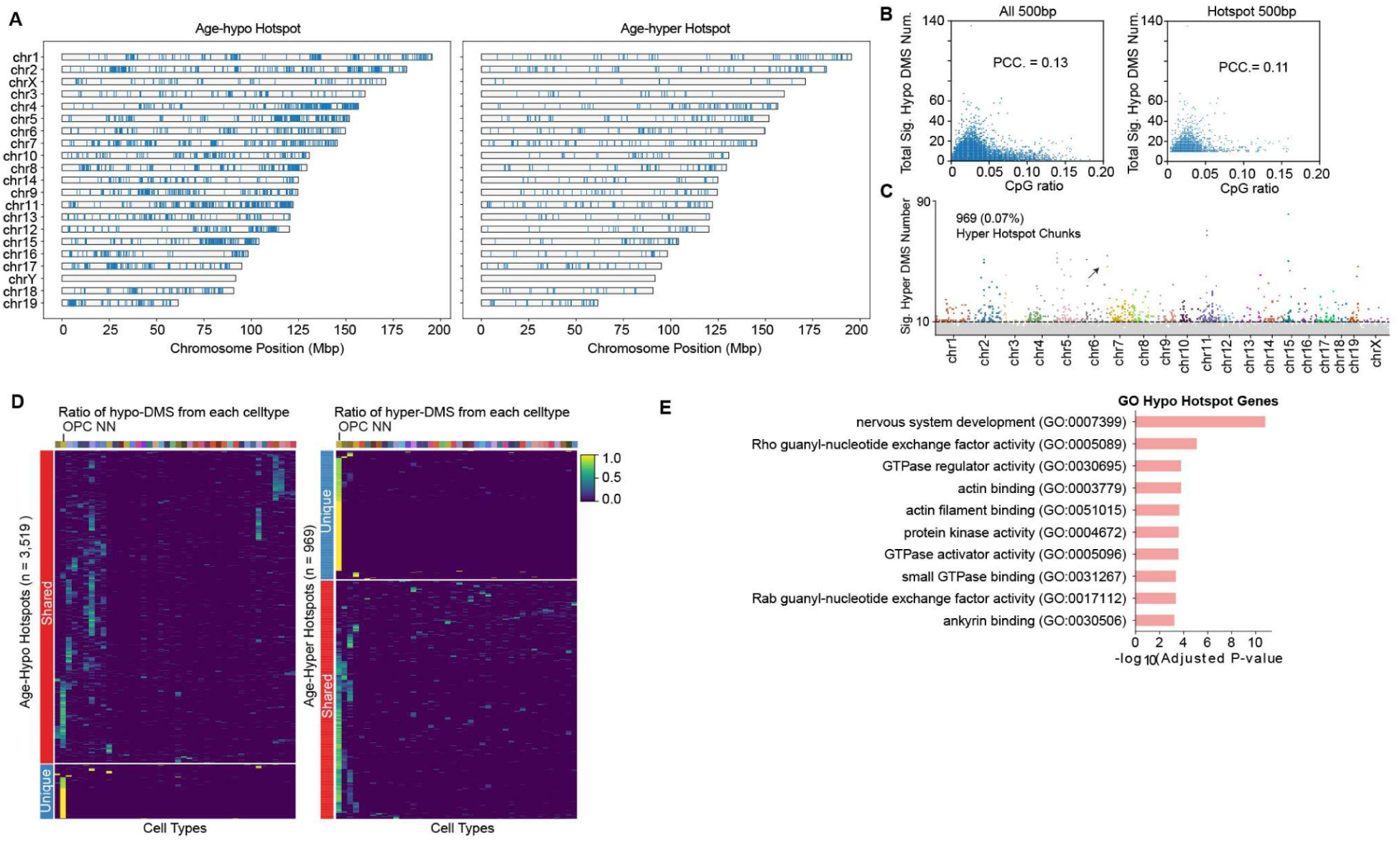
Age-related methylation hotspots. **(A)** Chromosome-wide visualization of age-hypo and age-hyper hotspot regions across the genome using a karyoplot representation **(B)** Scatterplot shows the relationship between CpG ratio and the total number of significant age-related DMS. Each dot represents a 500 bp chunk. **(C)** Manhattan plot shows the number of age-hyper DMSs (y-axis) on all 500bp genome chunks across chromosomes (x-axis). Each dot represents a 500 bp chunk colored by chromosomes. **(D)** Cell-type-specific contribution to age-related methylation hotspots. The heatmap shows the proportion of DMS originating from each cell type (columns) within identified hotspot regions (rows). Left and right show age-hypo and age-hyper hotspots, respectively. **(E)** Gene Ontology enrichment analysis showing significantly enriched biological processes among age-hypo hotspot genes.

**Figure S8.**
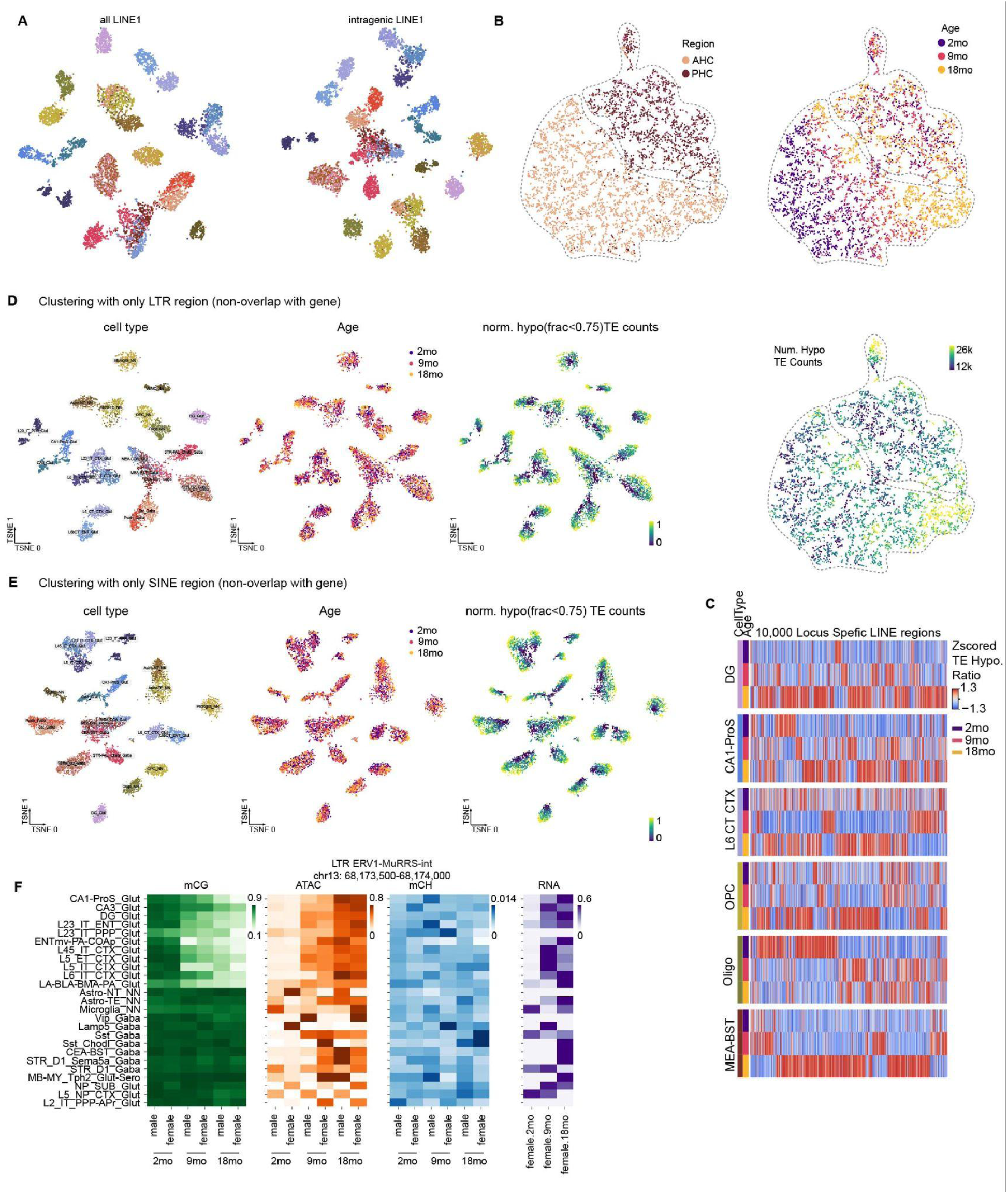
Clustering with retrotransposon regions. **(A)** t-SNE plots of cells clustered by LINE element regions (all regions, left; intragenic only, right), colored by cell type. **(B)** t-SNE of DG Glut neurons clustering based on intergenic LINE methylation, colored by brain region, age, and number of hypomethylated (mCG < 0.75) TE regions. **(C)** Normalized proportion of hypomethylated (mCG < 0.75) LINEs across age groups among cell types. **(D-E)** t-SNE of cells clustered by intergenic LTR **(D**) and SINE **(E)** regions, colored by cell type, age, and normalized hypomethylated region count (mCG < 0.75). **(F)** Multi-modal profiles of LTR elements by cell type (rows), showing mCG, ATAC, mCH, and RNA (left to right). Columns represent age/sex combinations.

**Figure S9.**
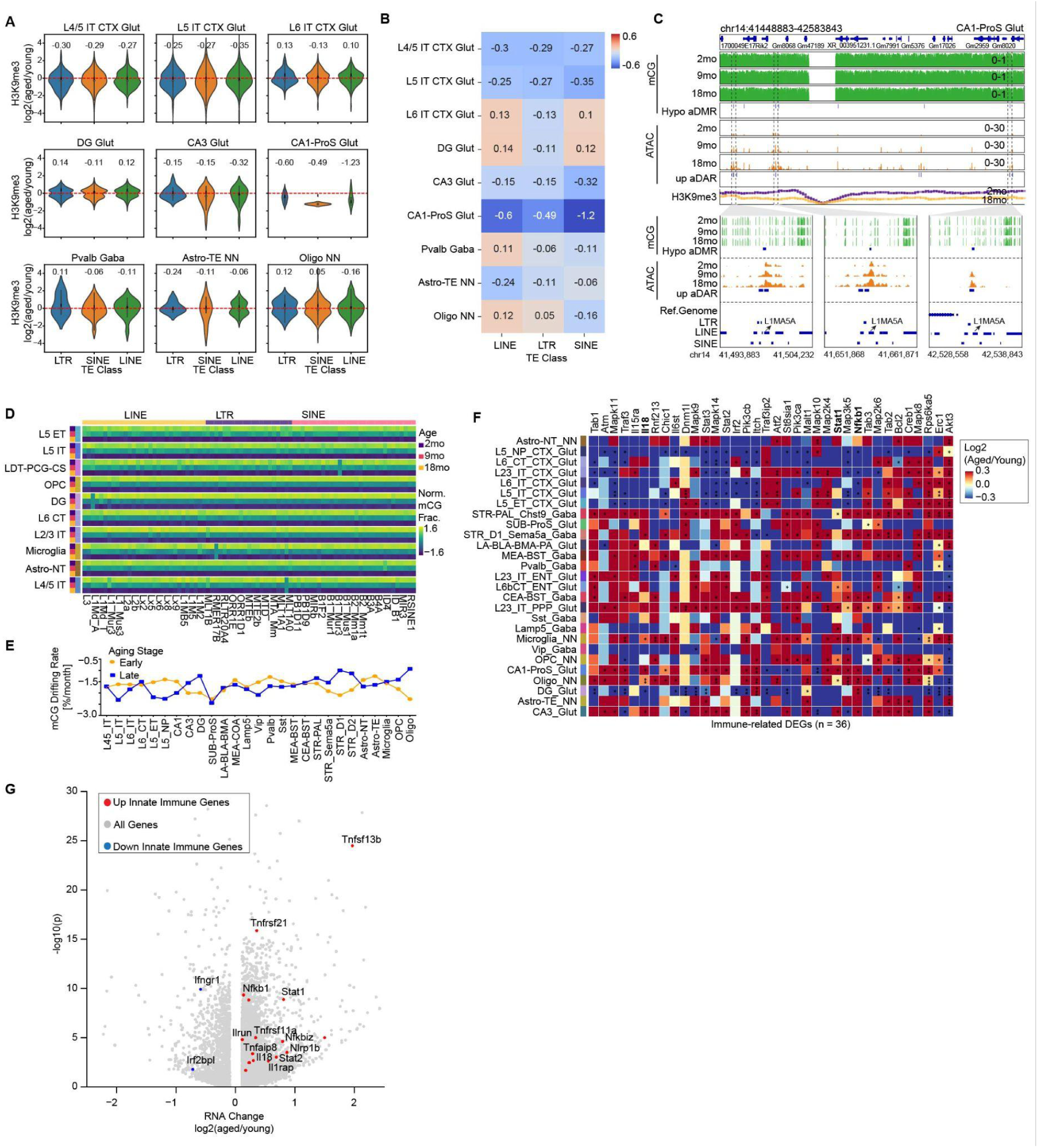
Age-associated TE derepression correlates with H3K9me3 loss. **(A)** Violin plots show the H3K9me3 (log2[18mo/2mo]) changes on hypomethylated TE regions across cell types. Numbers indicate average H3K9me3 change. **(B)** Heatmap representing average H3K9me3 changes (log2[18mo/2mo]) for distinct TE classes (columns) across cell types (rows). **(C)** Genome browser tracks showing epigenetic changes at chr14:41,448,883-42,583,843 in “CA1-ProS Glut”. (Top) mCG (green), age-hypo DMRs, ATAC (orange), age-up differential accessible regions (DAR), and H3K9me3 levels in young and old samples. Purple and orange represent 2-month and 18-month, respectively. (Bottom) Zoomed-in views of three different L1MA5A regions and their corresponding mCG and ATAC tracks. **(D)** Heatmaps showing normalized average DNA methylation levels of TE subfamilies across cell types and age groups. **(E)** Dot plot indicates the methylation drifting rate across cell types during early (2-month to 9-month) and late (9-month to 18-month) aging stages. **(F)** Heatmap depicting transcriptional changes in innate immune pathway genes across cell types. Color shows log2[18mo/2mo] changes and asterisks (*) indicate significance (FDR < 1e-3). **(G)** Scatterplot of age-DEGs in “Microglia NN”. Grey dots represent all age-DEGs; red and blue dots indicate upregulated and downregulated age-DEGs related to innate immune pathways.

**Figure S10.**
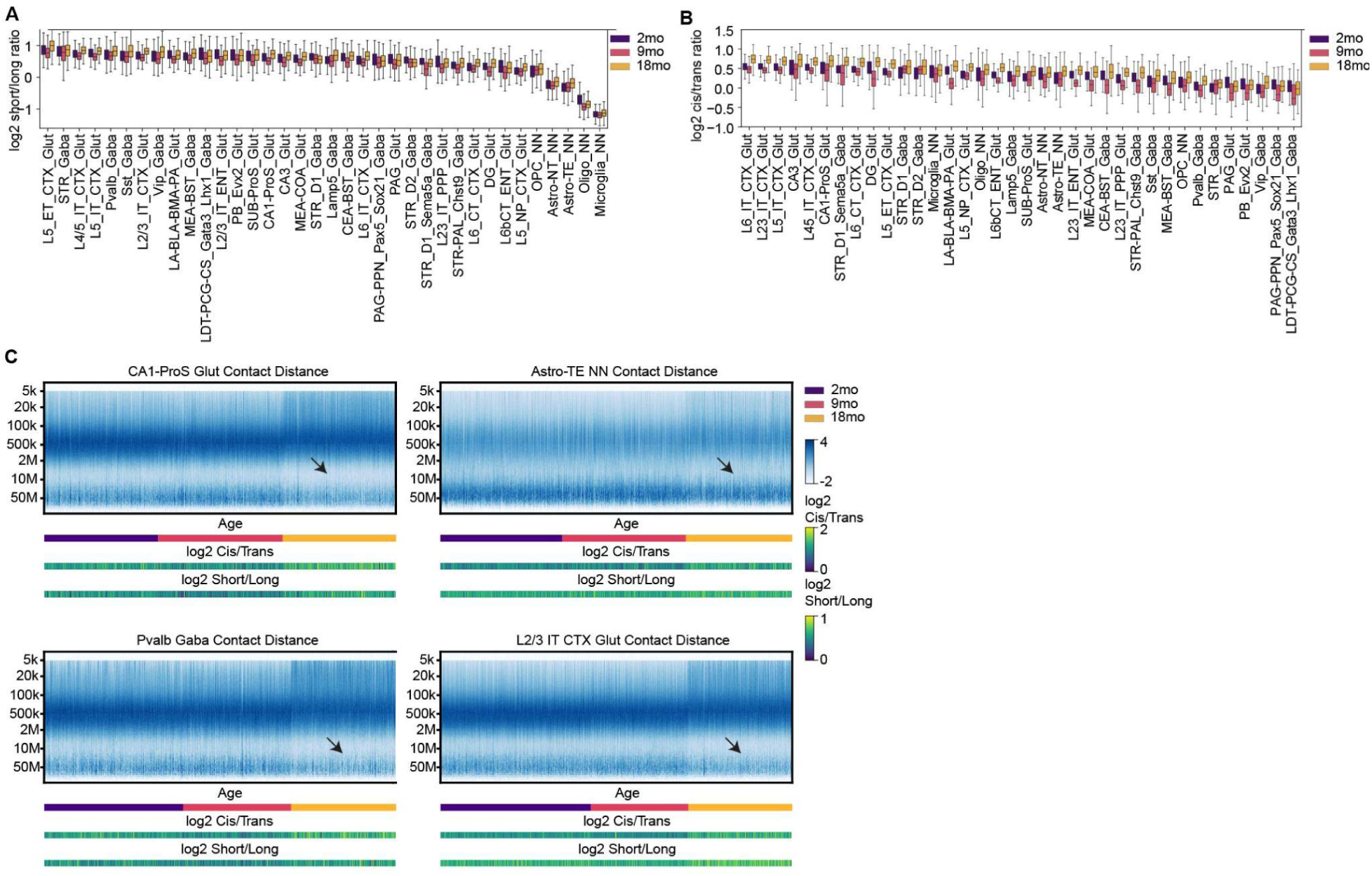
Age-dependent increase in short-to-long-range chromatin contacts. **(A-B)** Boxplot displays the **(A)** log2 ratio of short-range to long-range contacts and **(B)** log2 ratio of cis (intra-chromosomal) to trans (inter-chromosomal) contacts across age groups for each cell type. Cell types are ordered by the median ratio level. **(C)** Each panel is a cell type, including “CA1-ProS Glut”, “Astro-TE NN”, “Pvalb Gaba”, and “L2/3 IT CTX Glut”. Top: heatmaps showing frequency of contacts plotted against genomic distance in individual cells. Values are Z-score normalized within each cell (column). Cells are ordered by age groups. Y-axis shows genomic distance in log2 scale (Methods). Arrows indicate cells that show decreased long-range contact (10∼50Mb) ratios. Bottom: color bars displaying age, log2 short/long contact ratio, and log2 cis/trans contact ratio for each cell (column).

**Figure S11.**
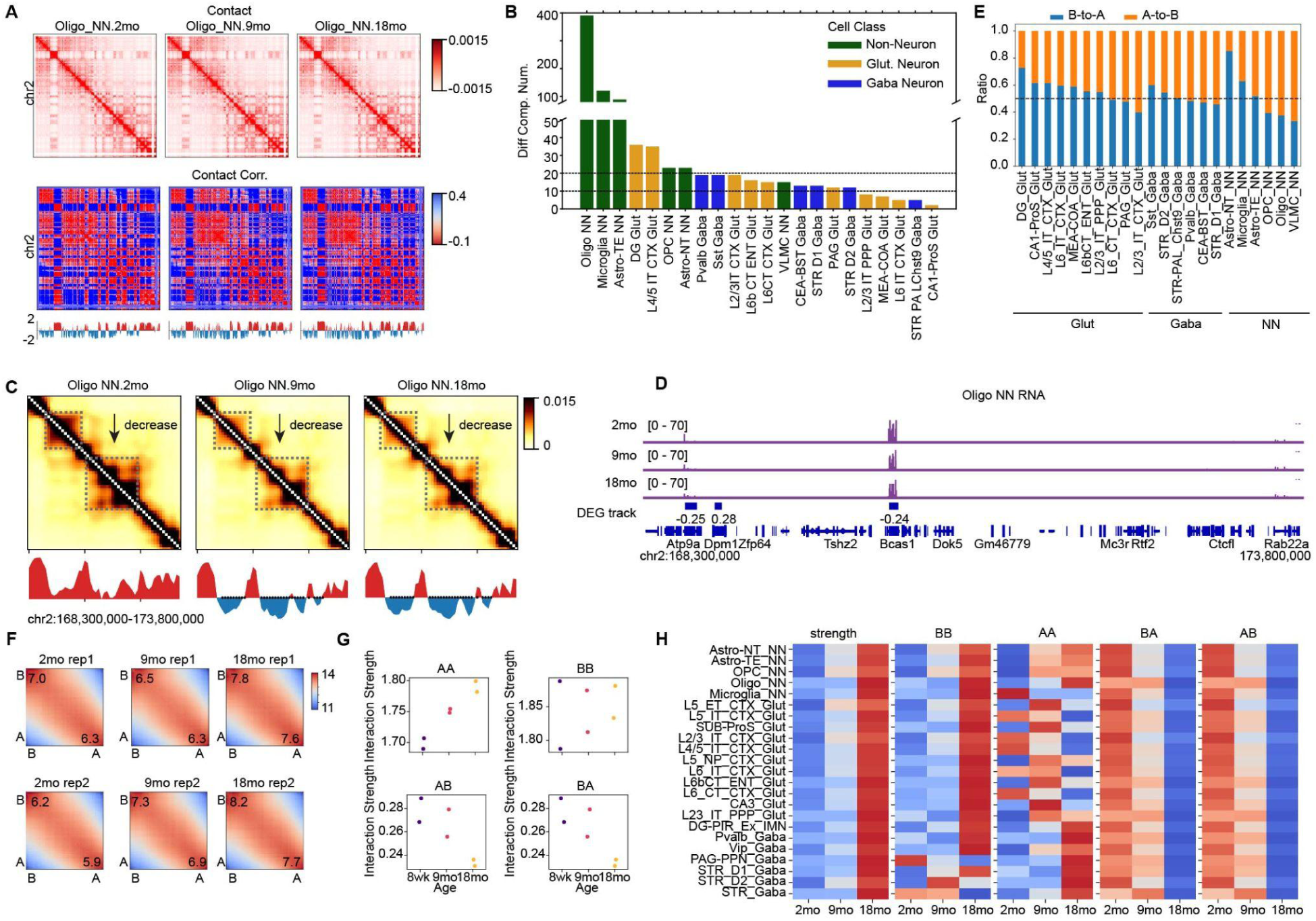
Age-dependent strengthening of chromatin compartmentalization and gene expression changes. **(A)** Top rows: Imputed contact maps of ‘Oligo NN’ among different age groups. Middle row: Heatmaps show the correlation matrices of distance-normalized contact maps in the top row, and Bottom rows: first principal component (PC1) analysis of correlation matrices, with positive (red) and negative (blue) values separated by zero baseline **(B)** Barplot shows the number of age-related differential compartments (age-DBs) across major cell types. Bars are colored by cell class. **(C)** Top: imputed contact maps of ‘Oligo NN’ among different age groups on chromosome 2, 168,300,000-173,800,000. Dashed squares highlight age-DÇs. Bottom line plots show the first principal component of the correlation matrices; red means the A compartment and blue means the B compartment. Black dots indicate age-differential compartment regions. **(D)** Genome browser displays RNA information of “Oligo NN” across age groups in the same region in **(C)**. **(E)** Barplot exhibits the ratio of B-to-A and A-to-B compartment switch across cell types, grouped by glutamatergic excitatory neuron (Glut), GABAergic inhibitory neuron (Gaba), and non-neuron (NN). **(F)** Saddle plots (Methods) of “Oligo NN” from different age groups and replicates. The axes are ranked by the compartment score of the group. Values are average distance-normalized raw contacts. The number at the corner represents the ratio between BB and BA interaction strength (top left) or the ratio between AA and AB interaction strength (bottom right). **(G)** Scatterplot shows the interaction strength between AA, BB, AB, and BA compartments across age groups in “Oligo NN”. Each dot is a replica colored by age. **(H)** Heatmap displays the Z-scored compartment strength (AA+BB)/(AB+BA), interaction strength between BB compartment, AA compartment, BA compartment and AB compartment among age groups across cell types (rows).

**Figure S12.**
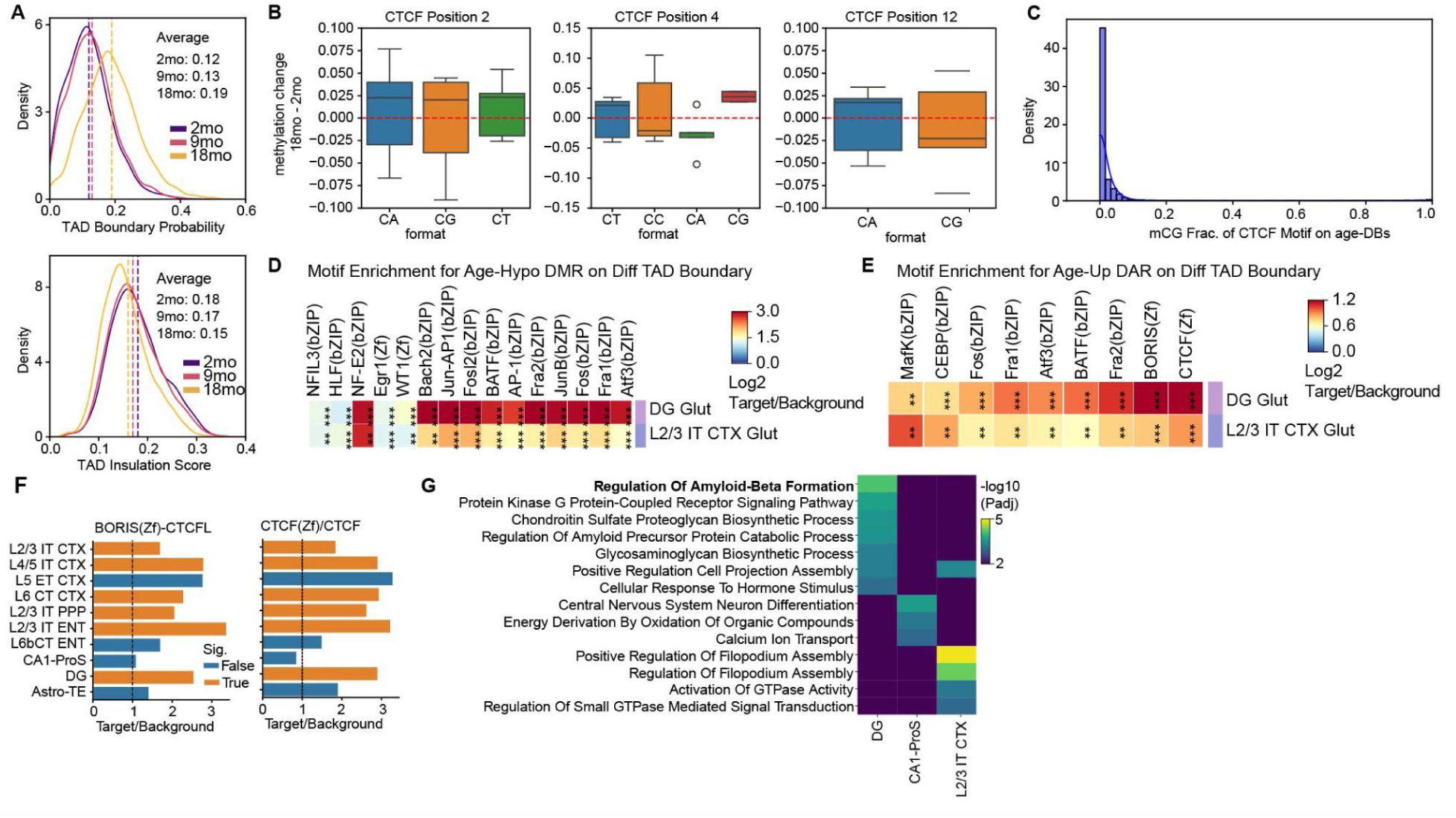
Age-related accessibility and methylation changes of CTCF binding sites. **(A)** Density plots illustrate the distribution of TAD boundary probability (top) and insulation score (bottom) across age groups for all age-DBs. Lines are colored by age. **(B)** Boxplot shows the methylation change of cytosines on CTCF peaks on age-DBs, grouped by their relation position on the CTCF motif. **(C)** Distribution of the mCG fraction level of CTCF sites on the age-DBs in the 2-month-old group. **(D-E)**, Motif enrichment of age-hypo DMRs **(D)** and age-up DARs on age-DBs **(E)** in “DG Glut” and “L2/3 IT CTX Glut” (FDR < 1e^-30^). “*” represents statistical significance and color represents log2(target/background). **(F)** Barplot shows the enrichment for CTCF motif across cell types. P-value < 0.05 is considered significant. **(G)** Heatmap shows the GO biological process enrichment for boundary-proximal genes across cell types (columns). Colors represent the statistical significance level.

**Figure S13.**
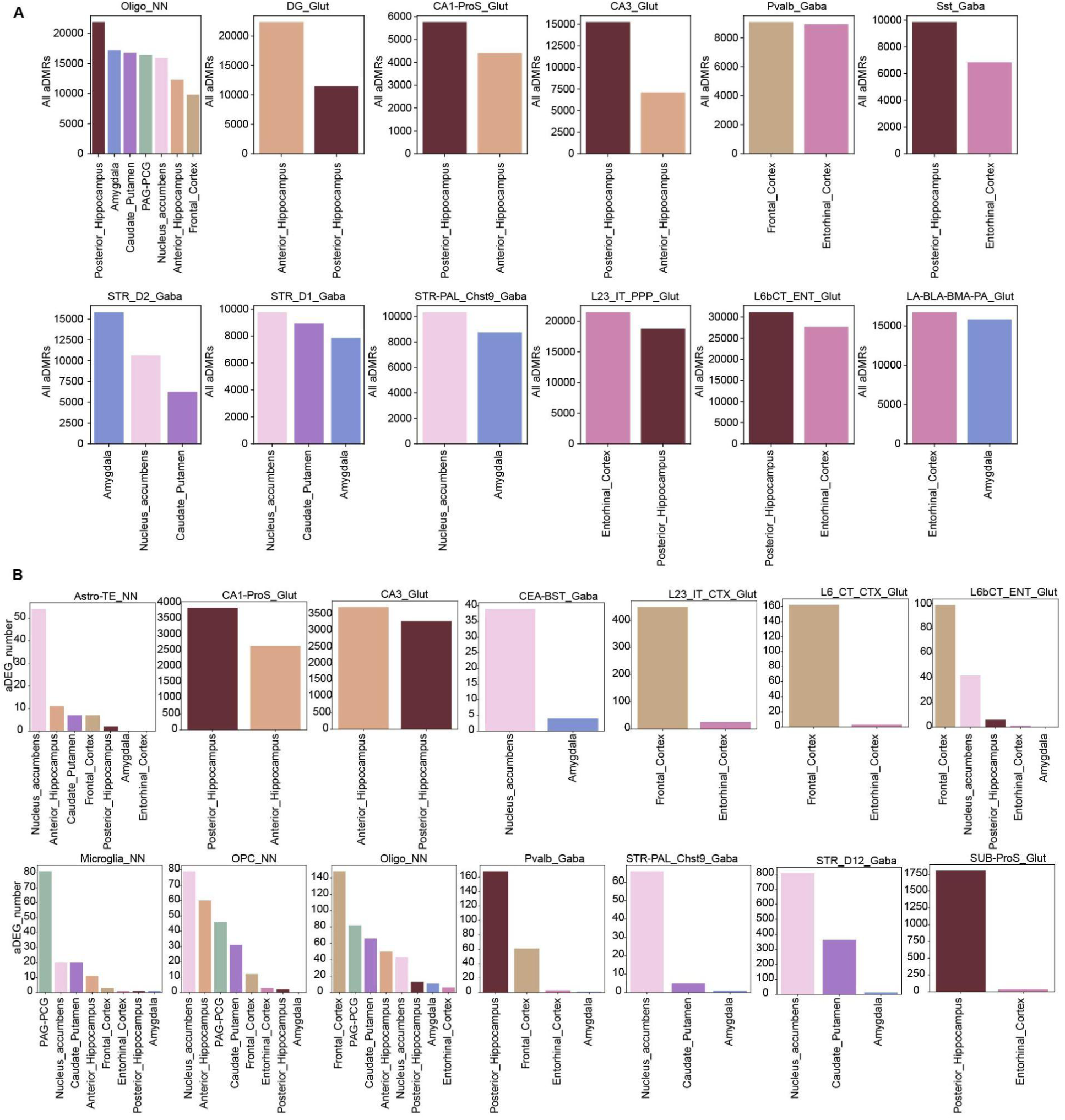
Regional heterogeneity of methylation and transcriptional changes. Number of region-specific age-DMRs **(A)** and age-DEGs **(B)** identified within each cell type after stratification by brain regions. Bars are colored by the brain regions.

**Figure S14.**
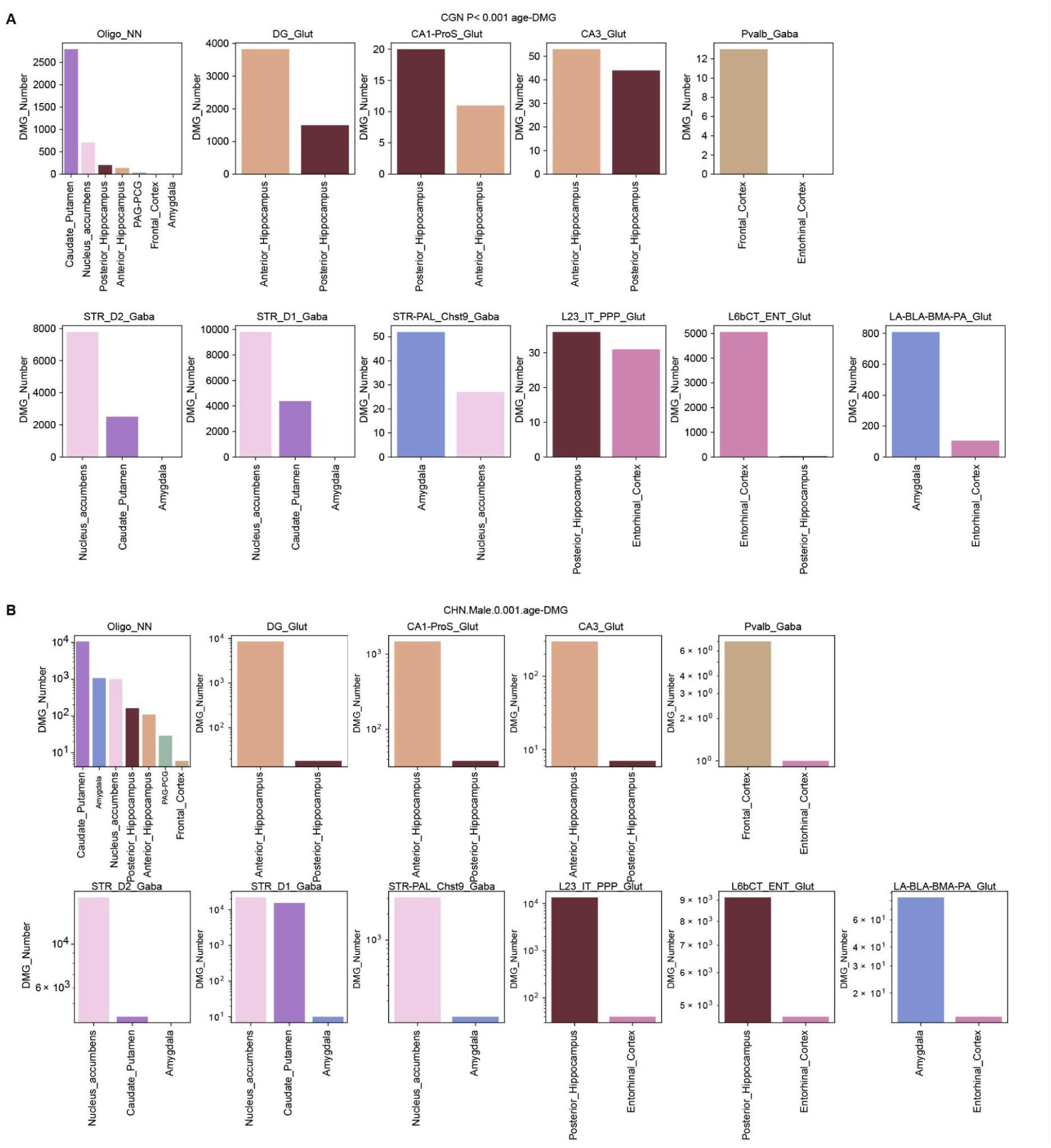
Regional heterogeneity of age-related gene methylation changes. **(A-B)** Number of region-specific age-related CH-DMGs **(A)** and CG-DMGs **(B)** identified within each cell type after stratification by brain regions. Bars are colored by brain region.

**Figure S15.**
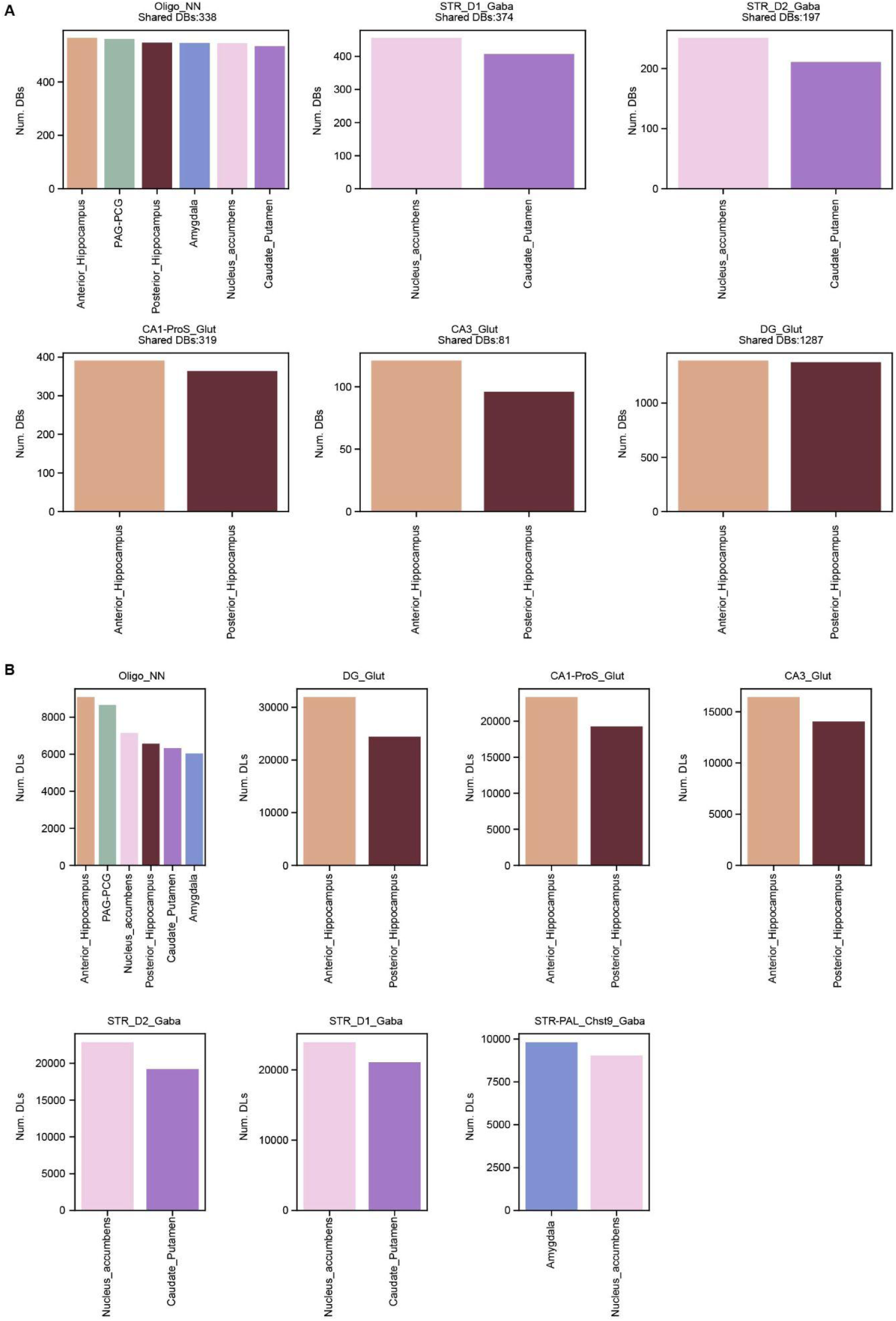
Regional heterogeneity of age-associated changes in TAD boundaries and chromatin loops. **(A-B)**, Number of region-specific age-associated differentially domain boundaries (age-DBs) **(A)** and differentially interacting loops (age-DLs) **(B)** identified within each cell type after stratification by brain regions. Bars are colored by brain region.

**Figure S16.**
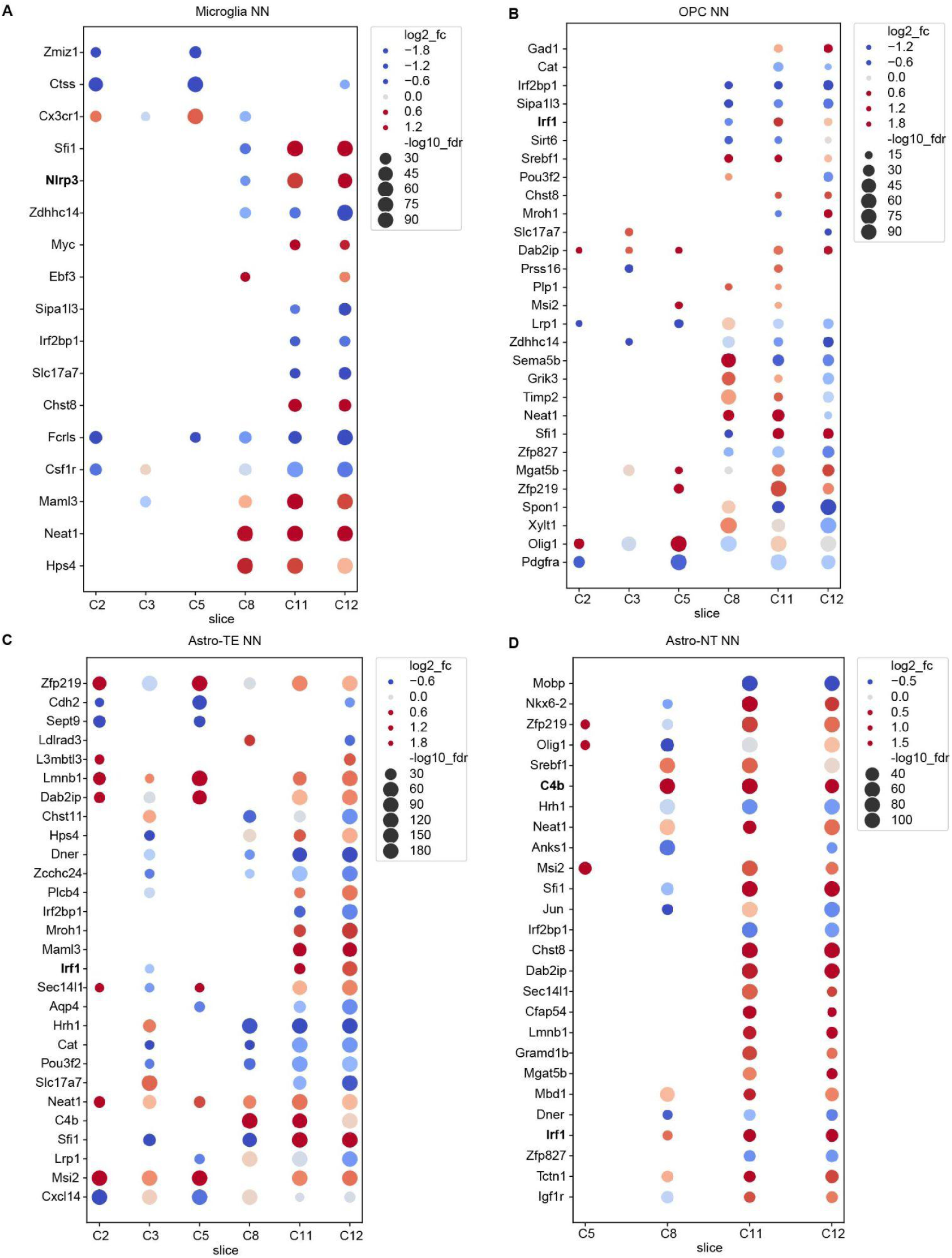
Anterior-to-posterior regional heterogeneity in age-related changes across glial cell populations. **(A-D)** Region-specific age-associated transcriptional changes in glia cells, including Microglia NN” **(A),** “OPC NN” **(B),** “Astro-TE NN” **(C),** “Astro-NT NN” **(D).** Columns represent MERFISH brain coronal slices ordered from anterior to posterior. Each dot represents a significant age-DEG within the specific MERFISH slice. Dot size indicates statistical significance (−log10[adjusted P]), and color intensity shows age-related fold changes (log2[18mo/2mo]).

**Figure S17.**
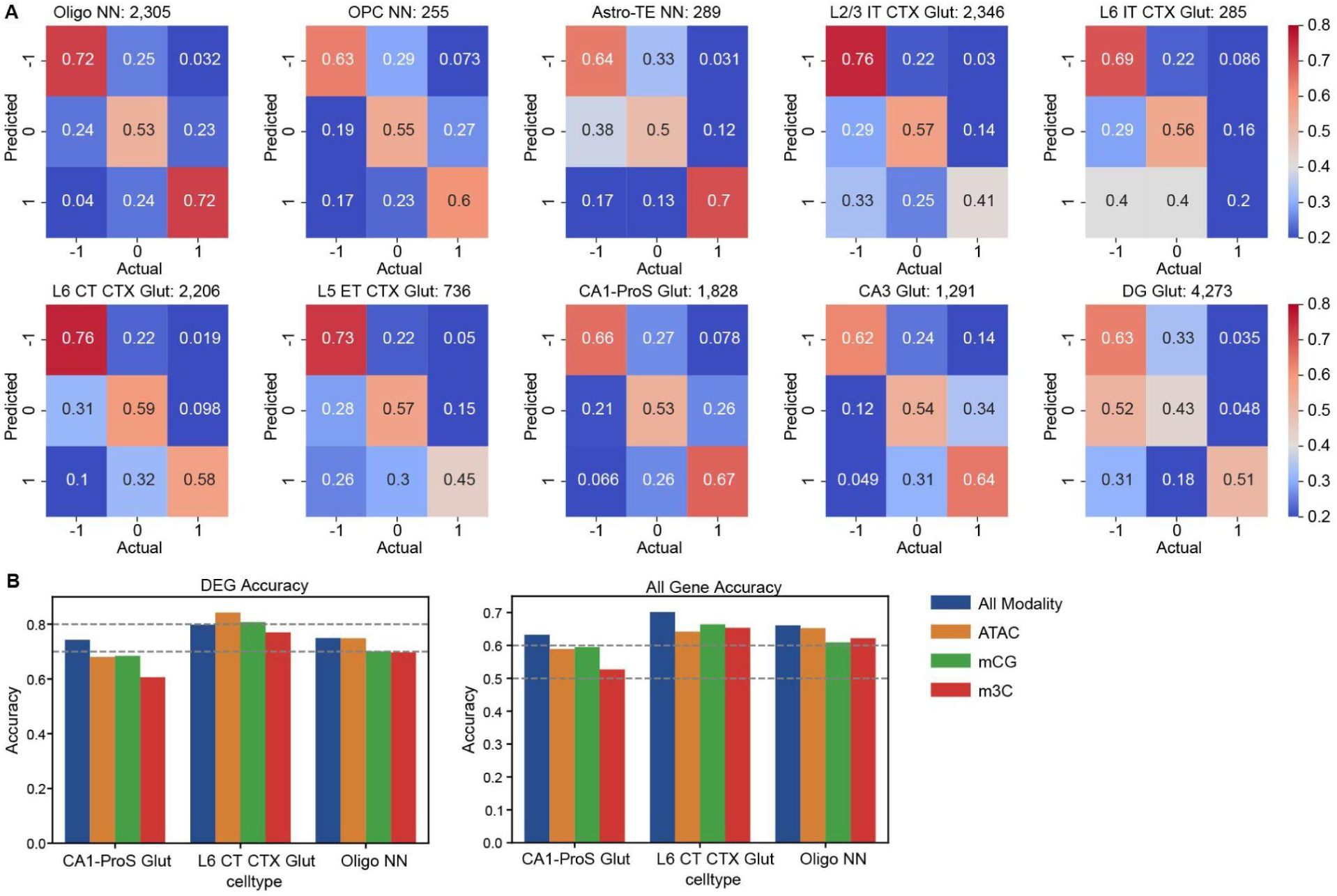
Multi-modal EAT model achieves higher prediction accuracy compared to single-modality approaches. **(A)** Confusion matrix showing prediction accuracy across cell types. Each column represents the true gene category, and each row represents the model-predicted gene category (−1: age-down DEG; 0: Non-DEG; 1:age-up DEG;). Color intensity and text indicate the accuracy of predictions for each gene category combination. **(B)** Barplot showing prediction accuracy using all modalities combined versus individual epigenetic modalities across three representative cell types: Oligo NN, CA1-ProS Glut, and L6 CT CTX Glut. The left panel depicts accuracy for age-DEGs, while the right panel shows accuracy for all genes.

**Figure S18.**
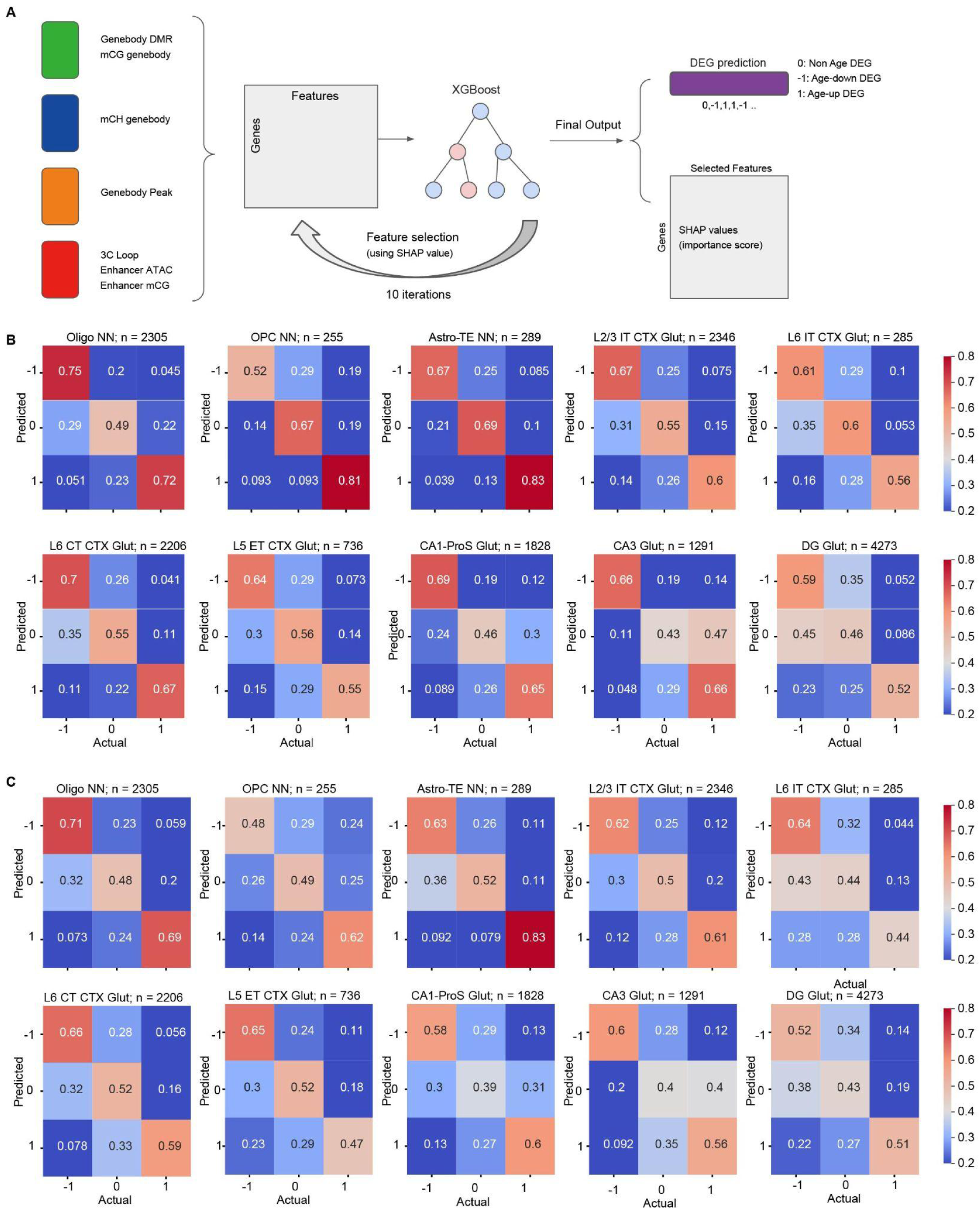
XGBoost model reliably predicts age-related transcriptome changes using epigenetic information. **(A)** Schematic representation of the XGBoost training and feature importance ranking workflow. **(B)** Confusion matrix showing prediction accuracy across cell types. Each column represents the true gene category, and each row represents the model-predicted gene category (−1: age-down DEG; 0: Non-DEG; 1:age-up DEG). Color intensity and text indicate the prediction accuracy for each gene category combination. **(C)** Same as **(B)** but using the Logistic Regression model for classification.

**Figure S19.**
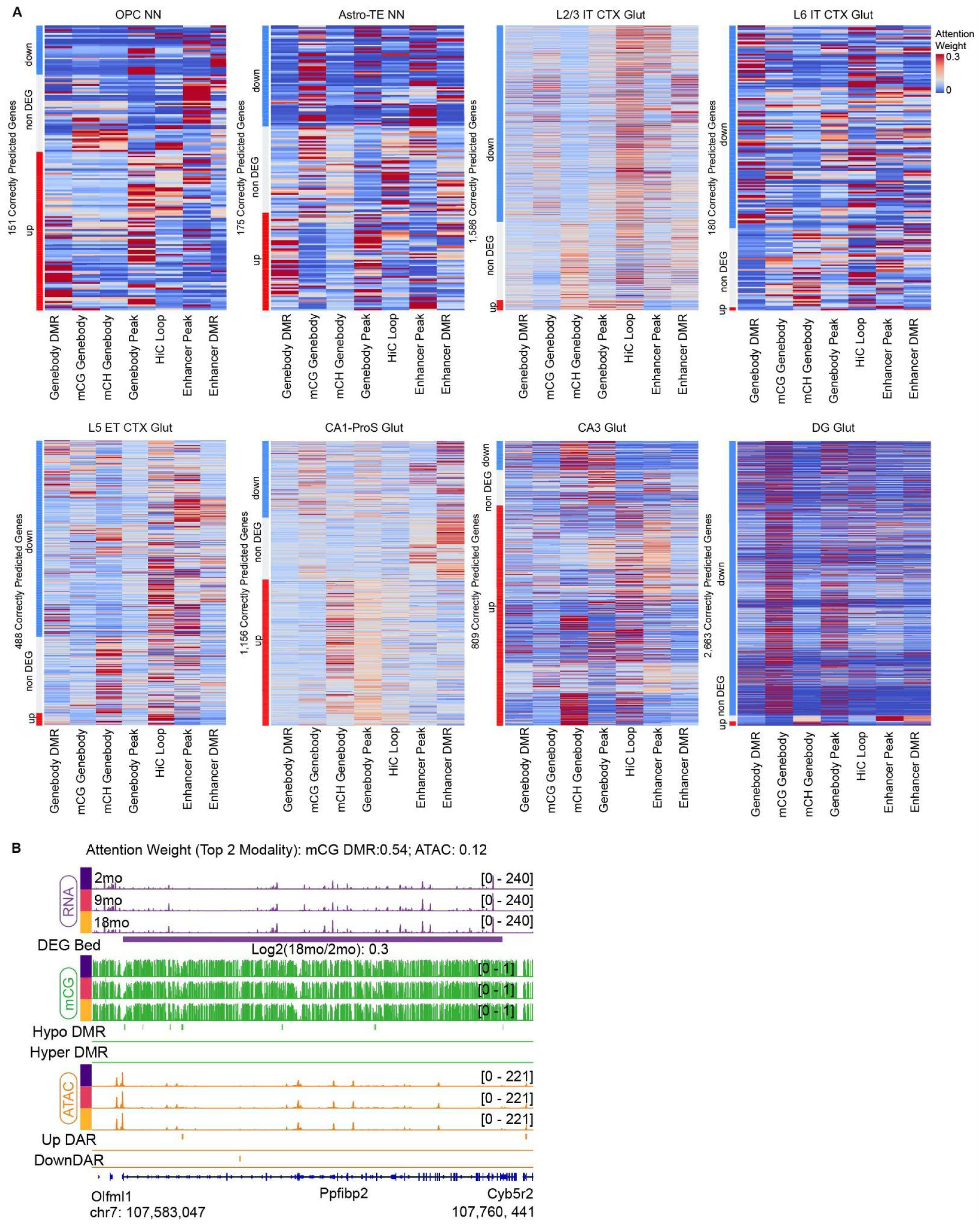
Contributions of epigenetic features in gene expression regulation. **(A)** Heatmap visualization of epigenetic-feature-specific attention weights for accurately predicted genes (rows) across cell types. Attention weights within each row sum to 1, with color intensity indicating the relative contribution of each feature to the prediction. **(B)** Pseudo-bulk profiles of *Pbfibp2* in “Oligo NN” across age groups, alongside the genome browser view showing RNA (purple), mCG (green), and ATAC (orange).

